# Nonsense-mediated decay constrains transposable element-driven evolutionary innovation

**DOI:** 10.64898/2026.10.22.760879

**Authors:** Tobias Plowman, George Kassiotis

## Abstract

Nonsense-mediated decay (NMD) degrades transcripts carrying premature termination codons, but its role in controlling transposable element-derived transcript diversity is unclear. We assembled an NMD-extended transcriptome (∼217,000 transcripts, ∼80% unannotated) from long-read RNA sequencing of cell lines after NMD inhibition. NMD-regulated transcripts were enriched in TE exonization, particularly of long terminal repeat (LTR) and long interspersed nuclear element (LINE) integrations, largely as alternative isoforms of annotated genes. Comparative genomics showed that NMD-silenced TEs were evolutionarily younger, more diverged, contained younger, less conserved splice sites, and more often encoded functionally distinct protein isoforms compared to TEs tuned by NMD or in stably expressed transcripts. NMD activity was elevated in lung cancer and associated with improved survival in lung squamous cell carcinoma. We further show that NMD-regulated primate-specific *Alu*-exonizing isoforms of *DDX49*, *NEIL3* and *SETDB1* produced stable proteins with distinct biological activity effects. NMD, thus, buffers TE-driven transcriptomic innovation, controlling the reservoir of evolutionary novelty.

## Introduction

Nonsense mediated decay (NMD) degrades mRNAs containing a premature termination codon (PTC) upstream of a spliced exon-exon junction (EEJ), precluding translation^1–3^. It relies on proteins within the exon-junction complex (EJC) deposited at the EEJ or factors recruited to the stalled ribosome, including eukaryotic initiation factor 4A-III (eIF4A3)^4^, up-frameshift protein 1 (UPF1) and suppressor with morphogenetic effect on genitalia proteins 6 (SMG6), and 7 (SMG7) – losing any compromises NMD efficacy^5–7^.

In human disease, NMD induces dystrophin loss in Duchenne muscular dystrophy after PTC mutations in the *DMD* gene^8^, is responsible for the loss of tumor suppressors in cancer^9^, and its inhibition has been explored in neoantigen generation^10,11^. NMD is also implicated in physiological gene dosing, for example stochastically recognizing upstream 5’ untranslated region (UTR) open reading frames (ORFs) and promoting degradation of the otherwise intact mRNA^12–14^. Overall, NMD represents a process employed across taxa to regulate mRNA translation.

Conversely, transposable elements (TEs) are major contributors to genetic disruption. Over evolutionary time, TEs have parasitized the genome, now representing up to half of human DNA^15,16^. The fate of TEs is a balance between loss through purifying selection and host co-option, several examples of which have been shown to influence human physiology^17–21^. Host mechanisms of TE suppression have also been described to control TE replication and expression, via transcriptional silencing by DNA methylation^22^, KRAB zinc-finger proteins (KZFPs)^23^, and the HUSH complex^24,25^, as well as splicing repression by RNA-binding proteins^26,27^.

Here, we investigate whether NMD represents a yet unexplored mechanism by which widespread TE-driven innovation is safeguarded. We leverage long-read RNA sequencing (RNAseq) to build a *de novo* transcriptome assembly under NMD blockade. We combine this with short-read RNAseq to demonstrate non-redundancy in NMD proteins, that NMD has a significant tendency to target transcripts overlapping TEs, and that NMD activity is altered in lung cancer. We use comparative evolutionary genomics approaches to show new transcript diversity is driven by TEs, and that NMD-silenced TEs represent a particularly young and non-conserved pool. Furthermore, we show specific examples of the physiological outcomes when NMD-controlled TE exonization events are expressed, identifying protein isoforms with distinct functions that emerged at different evolutionary timepoints.

## Results

### Transcriptome diversity as a function of NMD activity

To assess transcriptome regulation by NMD, we treated six human cell lines with DMSO (vehicle) or an inhibitor of eIF4A3 (eIF4A3-in-1). To validate inhibition, we measured the induction of NMD target-transcripts; an *L2a*-exonizing isoform of *BAG1* (*BAG1-L2a*)^28^ (Fig. 1a), the lncRNA *SNHG15*, and a long isoform of *TATDN2* (*TATDN2-L*) (Fig. S1a), initially identified by aligning a dataset of short-read RNAseq data in NMD-blocked HeLa cells^29^ to a prior pan-cancer transcriptome^30^. All were elevated in the cell lines upon inhibition (Fig. 1a lower panel; Fig. S1b).

**Figure 1.**
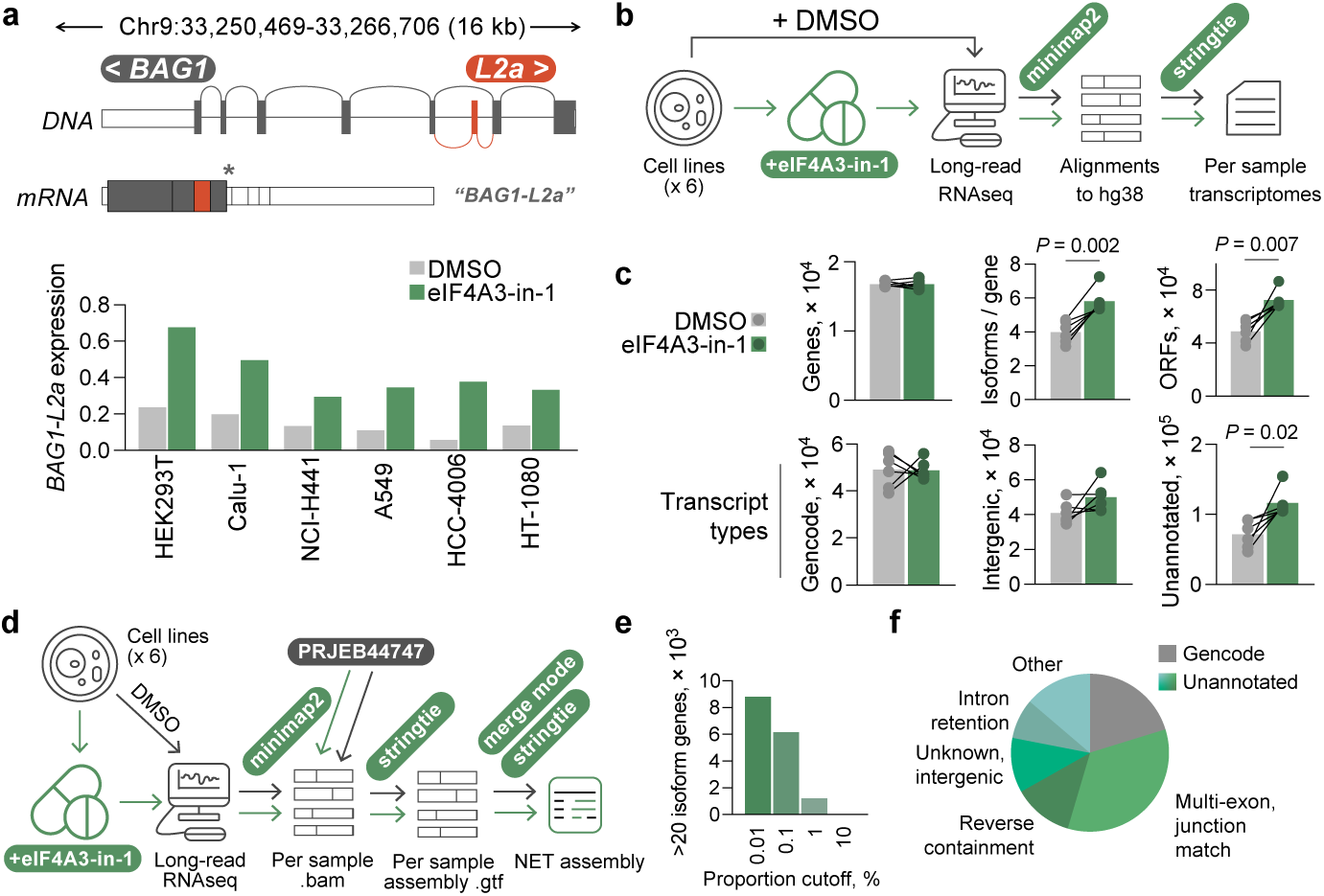
Generation of an NMD-Extended transcriptome (NET) assembly from long-read RNA sequencing of NMD-inhibited cell lines. **a**, Above: Schematic of the *BAG1* gene with alternative splicing to an *L2a* element to produce the *BAG1-L2a* transcript that encodes a PTC. Below: RT-qPCR of *BAG1-L2a* expression in cell lines (HEK293T, A-549, HCC4006, NCI-H441, Calu-1, HT-1080) treated with DMSO or 10 µM of eIF4A3-in-1 inhibitor for 24 h, relative to HPRT. **b**, Schematic of long-read RNAseq experimental and computational strategy to compare transcriptional diversity. **c**, Gene coverage, isoforms per gene, ORFs, and Gencode, intergenic, or unannotated transcripts derived from the long-read RNAseq of the 6 cell lines after vehicle (DMSO) or treated with 10 µM of eIF4A3-in-1 for 24 h. Symbols represent each cell line DMSO or eIF4A3-in-1 treated, and *P* values were calculated using a paired two-tailed Student’s *t* test. **d**, Schematic of strategy to generate the NET assembly. **e**, The number of genes with more than 20 isoforms, when varying the assembly strategy to include isoforms that represent a given proportion of the total locus. **f**, Transcriptional make-up of the NET assembly, derived from class codes annotated by Gffcompare.

We next long-read sequenced RNA from these samples, aligned reads to the hg38 genome reference and assembled them into transcriptomes using a custom pipeline (Fig. 1b). While the per sample transcriptomes had similar gene coverage, transcriptomic diversity was significantly elevated upon NMD inhibition; genes had more isoforms, the number of unannotated transcripts and predicted ORFs were significantly higher, whereas the number of intron-retaining, intergenic, and Gencode v49-annotated (Gencode) transcripts remained similar (Fig. 1c). The largest proportion of unannotated transcripts, and the most relative increase, were multi-exon transcripts that matched to one canonical splice junction (Fig. S1c).

To assess eIF4A3 blockade phenotype-specificity, we analyzed a long-read RNAseq dataset of HeLa cells^28^ treated with scramble siRNA (HeLa.control) or knockdown (KD) siRNA against UPF1, SMG6, SMG7, or both SMG6 and SMG7 (doubleKD). Here, only the doubleKD led to an increase in isoforms per gene and unannotated transcript numbers, with similar gene coverage and canonical transcripts as compared to HeLa.scramble (Fig. S1d). Overall, these results were consistent with a role for eIF4A3 and SMG6/SMG7^28^ in enabling NMD activity, and NMD in the regulation of transcript diversity.

Then, we combined the data from our eIF4A3-in-1 experiment with long-read HeLa cell data^28^ into an NMD-Extended Transcriptome (NET) assembly, employing quality metrics to determine optimal settings (Fig. 1d-e, S1e). Our NET assembly contained ∼217k transcripts, ∼44k (∼20%) matched Gencode, the remaining ∼80% unannotated were mostly exon junction-matching isoforms of genes (∼74k transcripts, 34%), although other minor classes were present (Fig. 1f).

### Differential sensitivity of an NMD-regulated transcriptome

We aligned and quantified the NMD-blocked HeLa short-read data^29^ using our NET assembly. The doubleKD had the most significant transcriptional effect and clustering revealed that each KD occupied a distinct region of t-SNE space (Fig. 2a). We then performed differential expression analysis on each KD condition, comparing with transcripts expressed in HeLa.control (Fig. S2a). HeLa.doubleKD showed the strongest effect, with 14,987 transcripts upregulated (log_2_FC > 1, *P*-adj < 0.05). UPF1 KD had a weaker transcriptional effect, while SMG6 and SMG7 KD had smaller effects (Fig. S2a).

**Figure 2.**
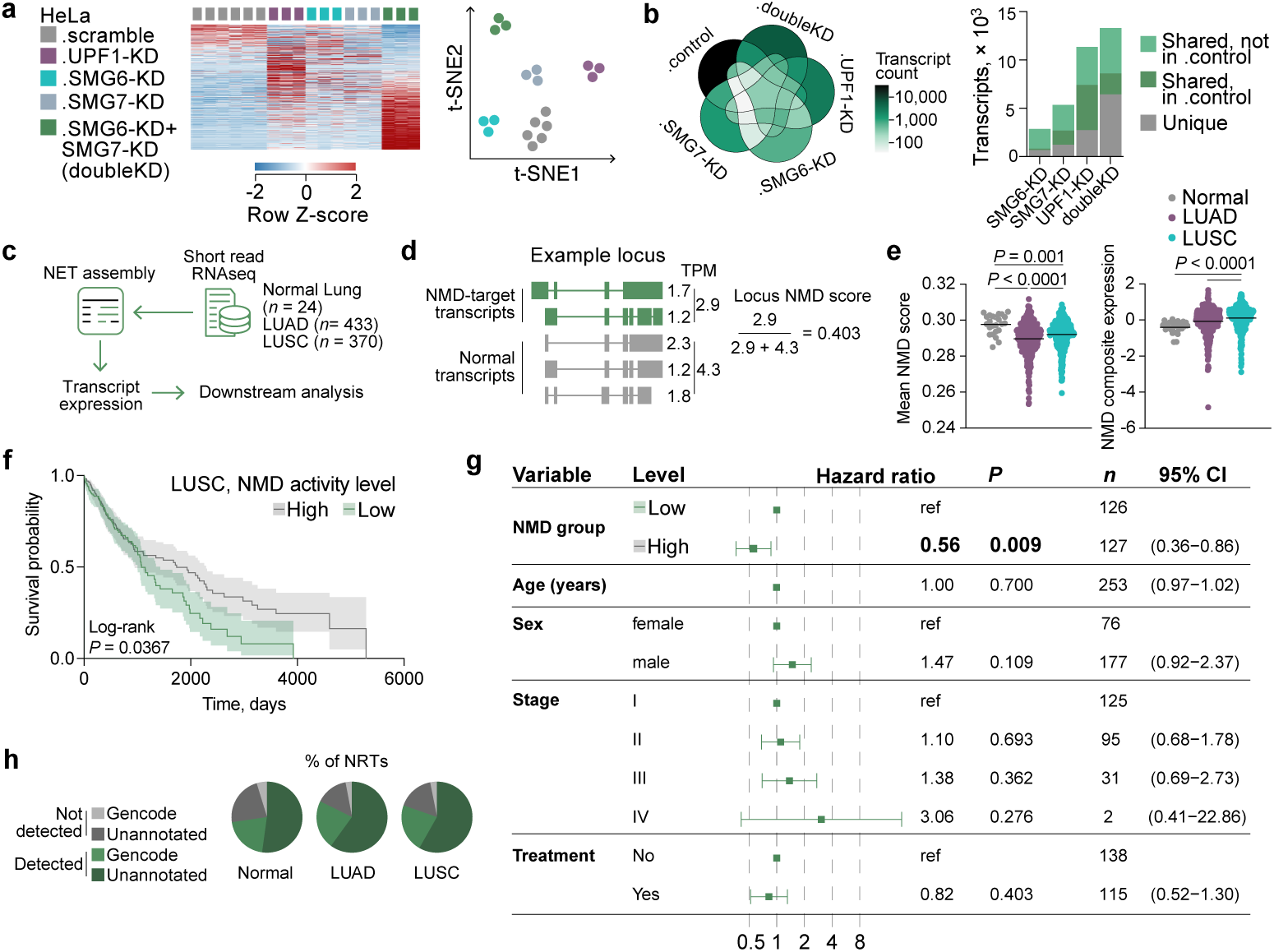
NMD pathway proteins are non-redundant, and NMD activity is altered in lung cancer. **a**, Left: Heatmap of transcripts significantly differentially regulated (≥2-fold change; *P* ≤ 0.05; *q* ≤ 0.05) after transfection with scramble siRNA or KD siRNA against NMD pathway proteins. Columns represent independent replicates for each condition. Right: t-SNE analysis of the same replicates and data, colored by condition. **b**, Left: Venn diagram overlap of the transcripts upregulated by each KD condition and compared to HeLa.control. Right: The number of transcripts upregulated by each KD condition, and colored by whether they overlap with another KD condition, and are expressed by HeLa.control. **c**, Schematic of NET assembly alignment and quantification using TCGA Normal Lung, LUAD, and LUSC samples. **d**, Schematic of method used to calculate per locus NMD score values that were averaged across a given patient to give an overall NMD activity score. **e**, Left: Mean NMD score per donor across the three cohorts. Each symbol represents an individual donor’s average NMD score across 6,408 commonly expressed loci. Right: NMD pathway composite expression score per donor. *P* values were calculated using a Kruskal-Wallis test followed by Dunn’s test for pairwise post-hoc comparisons with multiple-testing correction. **f**, Kaplan-Meier plot of overall survival probability in LUSC patients stratified into two groups by median NMD activity level. The *P* value was calculated with a log-rank Mantel-Cox test. **g**, Forest plot of a multivariate Cox proportional hazards model for overall survival in LUSC patients, with NMD activity group (low vs. high), age, sex, tumor stage (I-IV), and treatment status included as covariates. Squares indicate hazard ratios and error bars indicate 95% confidence intervals relative to each variable’s reference level. *n* indicates the number of patients in each group. **h**, Pie charts showing amongst the NRTs derived from the NET assembly, how many were detected in each cohort, and whether they were annotated or unannotated in Gencode.

6,432 (42.9%) of the 14,987 transcripts upregulated in HeLa.doubleKD were unique, and 6,889 (46.0%) were shared with one or more KD conditions, with 4,725 of the 6,889 not expressed in HeLa.control (Fig. 2b, S2b). Equivalent comparisons for the other KD conditions revealed that a higher proportion of upregulated transcripts were shared across KD conditions rather than being unique, such as the 75.8% of transcripts that were shared in UPF1-KD. Around half of the total transcripts upregulated and shared between multiple KD conditions were not expressed in HeLa.control (Fig. 2b, right).

These data therefore reflect the differing efficacy of NMD relative to the substrate transcript. They also demonstrate that while the loss of individual NMD pathway proteins can have some specific transcriptional effects, their functions broadly overlap in enabling NMD. 23,561 transcripts represented the union of those upregulated by any NMD KD, without removing any expressed by HeLa.control, defining NMD-regulated transcripts (NRTs).

### Higher resolution of the NMD-regulated lung cancer transcriptome

Given the suggested role for NMD in cancer and that our NET assembly was effectively an expanded NMD transcriptome for lung cancer (derived from lung cancer cell lines), we examined whether our NET assembly revealed changes in the NMD-regulated transcriptome in tumorigenesis using The Cancer Genome Atlas (TCGA) Lung Adenocarcinoma (LUAD), Lung Squamous Cell Carcinoma (LUSC), and adjacent normal lung tissue (Normal) data (Fig. 2c). Just under half of all transcripts in our NET assembly were expressed across the cohorts (Fig. S2c).

We then assigned an NMD activity score to each donor (methods) (Fig. 2d). Compared with Normal, both lung cancer cohorts exhibited a significant decrease in the mean NMD ratio, implying an increased level of NMD activity (Fig. 2e, left). In contrast, tumor stage, sex, age at diagnosis and treatment status did not correlate with changes in the mean NMD ratio (Fig. S2d-g).

The change in NMD ratio associated both with a significant increase in non-NRT expression, as well as a decrease in NRT expression (Fig. S2h). Moreover, there was a significant increase in overall expression of NMD pathway proteins in lung cancer, measured by an NMD composite score (Fig. 2e, right), which aggregated expression for a set of core NMD proteins, many of which were significantly overexpressed in lung cancer individually (Fig. S2i).

Lower NMD activity (corresponding to a high NMD ratio) correlated with lower overall survival probability in LUSC (median 1075 vs. 1736 days; log*-*rank *P* = 0.0367; HR = 1.41, 95% CI 1.02-1.95, Mantel-Haenszel) (Fig. 2f), with the divergence occurring predominantly at later follow-up timepoints (Gehan-Breslow-Wilcoxon *P* = 0.42). This association remained significant in a multivariate analysis (High NMD activity HR = 0.56, *P* = 0.009) (Fig. 2g) and, of all the variates tested here (age, gender, stage, treatment status), NMD activity was the only that associated significantly with survival. The trend was the opposite in LUAD, lower NMD activity was protective (Fig. S2j), however this was not significant in a multivariate analysis (HR = 1.4, P = 0.088) (Fig. S2k), where stage and treatment status were instead significant covariates.

The majority of NRTs detected in the three cohorts were newly identified in our NET assembly (Fig. 2h). Thus, these data demonstrated the increased resolving power provided by our *de novo* assembled NET in identifying NMD-related transcriptomic changes in cancer and that NRTs readily expressed in cancer tissue may have clinical significance.

### NRTs are enriched in TEs

We hypothesized that NMD-regulated transcriptomic diversity could derive from TE exonization events, given our and others’ previous efforts to explore the overlooked TE-driven transcriptome^17–20^. NMD inhibition can have indirect effects, so we predicted direct targets in our NET assembly using TranSuite^31^. 60,831 (28%) direct transcript targets overlapped 9,107 (39%) of the 23,561 NRTs (Fig. 3a, S3a), defining TransSuite-filtered NRTs (tsNRTs).

**Figure 3.**
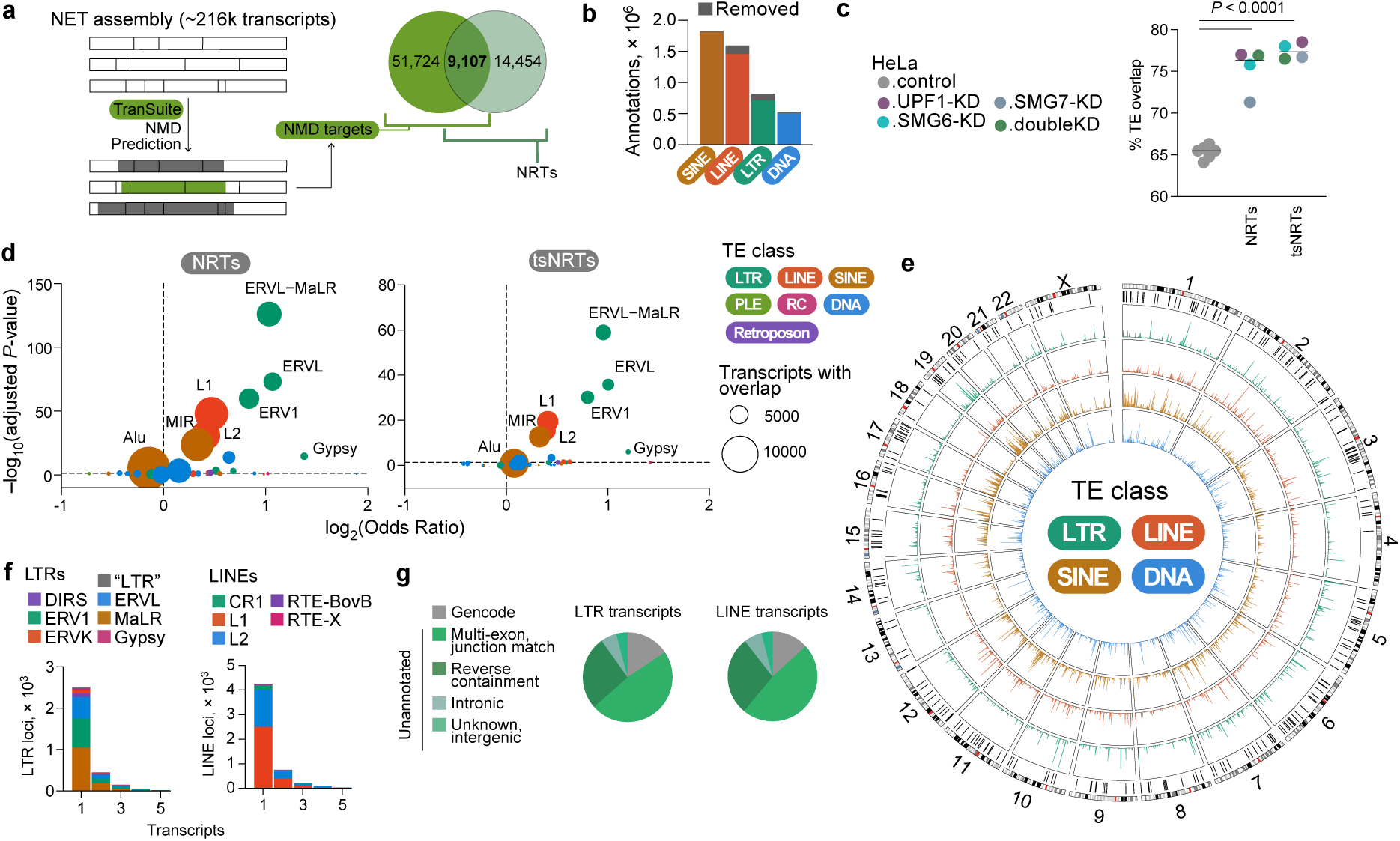
NMD has a significant tendency to regulate TE-overlapping transcripts of known genes, especially LTR and LINE elements. **a**, Schematic showing TranSuite NMD prediction in the NET assembly, and the transcript overlap between those and the NRTs. **b**, The number of and reduction in TE annotations per class after modifying the raw RepeatMasker/Dfam3.9 hg38 annotation. **c**, The proportion of transcripts overlapping a TE within HeLa.control, or those significantly upregulated by each of the NMD KDs, without (NRTs) or with TranSuite prediction overlap (tsNRTs). *P* values were calculated with one-way ANOVA. **d**, TE family enrichment among TEs overlapping NRTs (left) and tsNRTs (right) relative to TE-overlapping transcripts in HeLa.control. Each point represents one TE family, point color indicates TE class, and point size indicates the number of transcripts with that TE family overlap in the NRTs (left) or tsNRTs (right). Odds ratios and adjusted *P* values were calculated using a two-sided Fisher’s exact test with BH correction. The dashed horizontal line marks adjusted *P* = 0.05, and the dashed vertical line marks an odds ratio of 1. **e**, Circos plot of the chromosomal distribution of TE-class overlaps in NMD-TEs, with each TE class (LTR, LINE, SINE, DNA) shown as a separate track and colored as indicated. Black vertical lines on the second-most outer ring indicate 1-Mb genomic windows significantly enriched in NRT-derived TE loci relative to HeLa.control TE loci by one-sided Fisher’s exact test (BH-adjusted *P* < 0.05, odds ratio > 1). **f**, The number of NRTs overlapped by LTR (left) and LINE (right) loci, colored by subfamily. **g**, Pie charts of the type of NRTs overlapped by LTR (left) and LINE (right) elements, derived from the NET assembly Gffcompare annotation.

To compare TE overlap, we annotated TEs in the human genome. We employed a custom pipeline to remedy annotation fragmentation (Fig. S3b-c), which reduced total TE annotations (Fig. 3b), and these were intersected with NRTs/tsNRTs and HeLa.control transcripts. There was a significant tendency for NRTs/tsNRTs to overlap a TE compared to HeLa.control transcripts (Fig. 3c). We tested for transcript length as a confounder, but comparing a matched-length sample of HeLa.control transcripts reproduced the same TE overlap propensity (Fig. S3d). In the NRTs and tsNRTs compared to HeLa.control, there was significant enrichment in the major LTR families (ERV1, ERVL-MaLR, ERVL) and LINE elements (L1, L2) (Fig. 3d). This enrichment was not driven by any one KD, as comparing each individually revealed the same enrichment profile (Fig. S3e).

As orthogonal exploration, we aligned and quantified short-read RNAseq data from the HT1080 cell line after SMG7 knockout (KO)^32^ and performed differential expression analysis (Fig. S3f). There was significant enrichment in TE overlap (Fig. S3g), especially LTR and LINE elements (Fig. S3j), in transcripts upregulated in SMG7 KO compared to HT1080.control.

The TEs exonized in tsNRTs (NMD-TEs) were distributed around the genome, but there were 258 one-megabase windows with selective NRT production (Fig. 3e, outer layer). However, at the individual transcript level almost all were single TE copies exonized into one transcript (Fig. 3f). Initially, we wondered whether intact TEs may be subject to NMD regulation, as splicing in LTR elements is known to occur^33,34^. However, the LTR-and LINE-overlapping transcripts mapped mostly to Gencode, or to unannotated isoforms of known genes (Fig. 3g). The remaining transcripts mapped to intergenic or unknown regions that were in fact hg38 scaffolds or loci producing unannotated chimeras of TE and non-TE genomic regions (Fig. S3j).

Altogether, these data suggested that the tendency of NMD to regulate LTR and LINE elements observed here was a result of their enhanced capacity to act as segments that increase transcriptomic diversity from annotated genes. Nevertheless, there were also cases of transcript production from loci not previously described to be transcriptionally active, revealed by our NET assembly.

### NMD targets younger TEs

To understand the evolutionary context of NMD-regulated TEs, we performed a comparative genomics and conservation analysis in ten species, using four TE groups; i) a randomly sampled set of 50k TE loci matched to TE class proportions in the human genome (Genomic-TEs), ii) TEs expressed by HeLa.control and not among tsNRTs (Stable-TEs), iii) TEs in tsNRTs expressed by HeLa.control (Tuned-TEs), and iv) TEs in tsNRTs only detectable after NMD blockade in HeLa cells (Silenced-TEs) (Fig. 4a). TEs that only aligned to the chimpanzee genome were the youngest, suggesting their integration within the last six million years (Ma). TEs that aligned across all species were defined as integrations over 95 Ma old. Eight other genomes covered various speciation events between those two timepoints.

**Figure 4.**
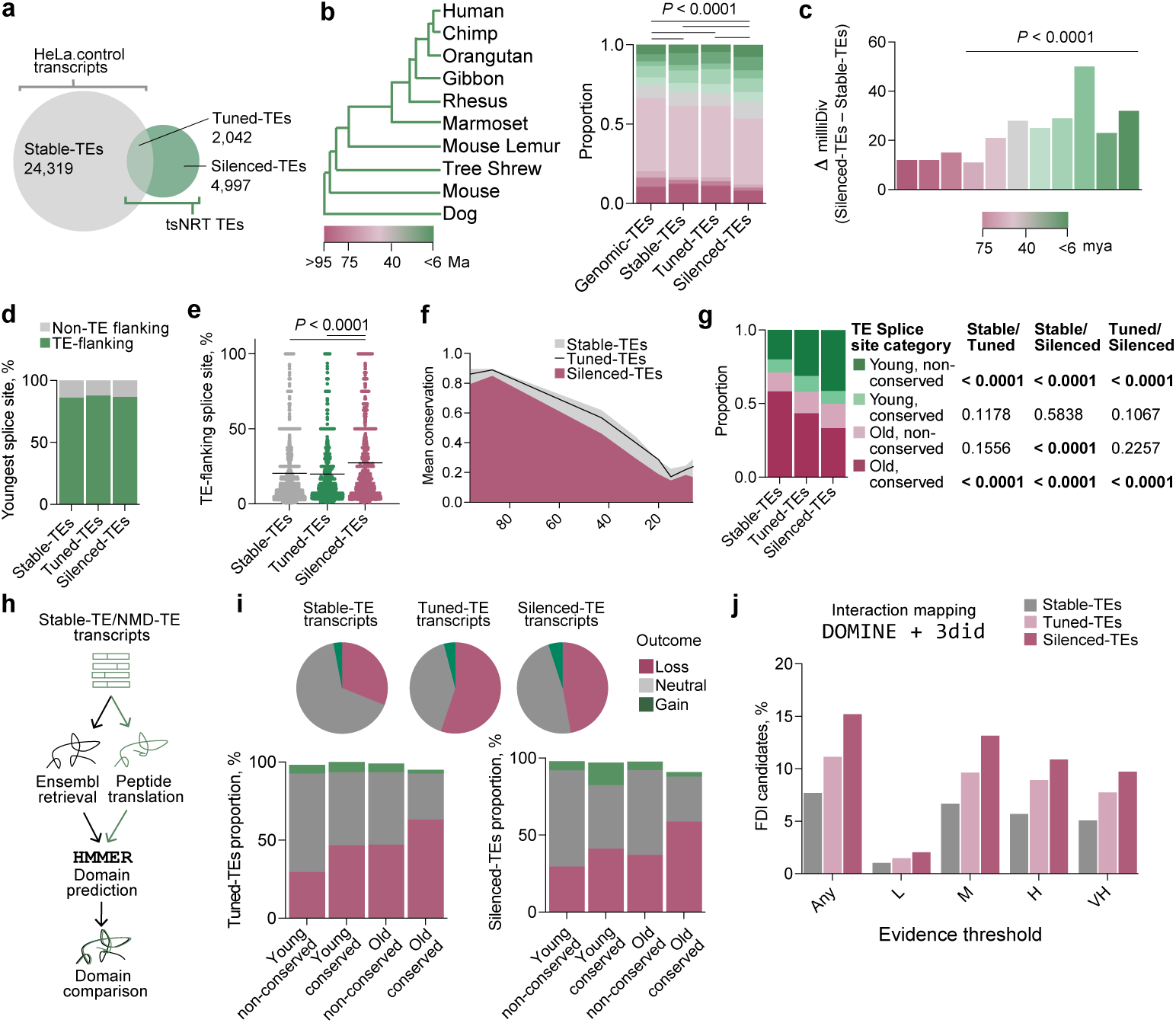
NMD regulates younger and less-conserved TEs that encode possible Functionally Distinct Isoforms (FDIs). **a**, Venn diagram of the three expressed TE transcript groups defined by HeLa.control and NRT TEs (NMD-TEs). **b**, Left: Evolutionary time tree of ten genomes used for TE integration age estimation, with color scale indicating age in million years (Ma). Right: The proportion of TEs within each age bin. *P* values were calculated with a chi-square test on the binned counts. **c**, The difference in milliDiv values between Silenced-TEs and Stable-TEs for the most diverged TE milliDiv per transcript (in case of overlapping multiple TEs). *P* values were calculated within each age bin using a two-sided Wilcoxon rank-sum (Mann-Whitney U) test. **d**, The proportion of transcripts whose youngest splice site flank or do not flank a TE. **e**, The proportion of splice sites that flank a TE per transcript. Each symbol represents an individual transcript, the line represents the median, and *P* values were calculated using a two-sided Wilcoxon rank-sum (Mann-Whitney U) test. **f**, Mean phastCons100way conservation score per transcript for TE-flanking splice sites, binned by the integration age of each transcript’s youngest TE-flanking splice site, derived from the liftOver splice site analysis. Human-specific splice sites excluded. **g**, The proportion of transcripts by the age of their youngest TE-flanking splice site (Old: ≥43 Ma; Young: <43 Ma), and their mean phastCons100way conservation score (conserved: ≥0.5; non-conserved: <0.5). Human-specific splice sites excluded. *P* values were calculated between groups within each category using a chi-square test with BH correction. **h**, Schematic of the pipeline used to identify and contrast domains present in TE transcript-encoded proteins relative to their canonical counterparts. **i**, Above: Pie charts of the domain change outcomes in TE transcript-encoded proteins relative to their canonical counterparts. Below: Stacked bar plots of the domain change outcomes in Tuned-TE and Silenced-TE transcript-encoded proteins relative to their canonical counterparts, binned by their youngest TE splice site category. **j**, The proportion of TE transcripts that encode possible FDIs, based on retention of an interacting domain combined with loss of an interacting domain, derived from the DOMINE interaction prediction database (L, low confidence; M, medium confidence; H, high confidence; VH, very high confidence). ‘Any’ is the overlap from the four predicted lists.

The set of genomes used encapsulated most TE ages – approximately 84% of human Genomic-TEs integrated between 6 and 95 Ma (Fig. 4b). Expressed TEs (Stable-TEs, Tuned-TEs, and Silenced-TEs) were significantly younger than Genomic-TEs (Fig. 4b). Silenced-TEs were significantly younger than all other TE groups in the comparison, while Stable-TEs and Tuned-TEs were not comparably different in age. This was not driven by any one TE family, there was significant enrichment in younger integrations from six TE families – the other three families were younger but not to statistical significance (Fig. S4a, left). Again, no families in Tuned-TEs were significantly younger than Stable-TEs (Fig. S4a, right).

We assessed TE divergence using milliDiv values (higher values indicate higher divergence), commonly used as a proxy for age, but may also depend on tolerated mutation rate and selection. For all age bin-matched TEs less than 75 Ma old, the median milliDiv was significantly higher in Silenced-TEs compared to Stable-TEs (Fig. 4c), but while Tuned-TEs had mostly higher median milliDiv values, none were significant (Fig. S4b). Altogether, this would suggest that Silenced-TEs are under weaker forces of purifying selection relative to Stable-TEs – they are tolerant to mutation or indeed were mutated leading to a frameshift/PTC introduction by purifying selection, in turn causing NMD targeting.

In 86.0%, 87.7%, and 86.7% of Stable-TE, Tuned-TE and Silenced-TE transcripts respectively, the youngest splice site flanked a TE-containing exon, suggesting TE exonisation is a key driver of diversity (Fig. 4d). Furthermore, splice sites in Silenced-TE and Tuned-TE transcripts were significantly younger than Stable-TEs, although this effect was subtle (Fig. S4c). Examining TE contributions to splicing, Silenced-TEs had a significantly higher average proportion of splice sites flanking a TE compared to Tuned-TEs and Stable-TEs (Fig. 4e). Moreover, the conservation of Silenced-TE splice sites was lower compared to Tuned-TEs and Stable-TEs (Fig. 4f). Dividing splice sites according to the age closest the median (43 Ma) revealed that across all transcripts, splice sites in Silenced-TEs were the youngest and the least conserved (Fig. 4g). Stable-TE splice sites were the oldest and most conserved and Tuned-TEs represented an intermediate (Fig. 4g). There was a significant tendency for Tuned-TE and Silenced-TE transcripts, but not Stable-TE transcripts, to involve younger genes compared to all genes expressed by HeLa.control (Fig. S4e). However, Tuned-TE and Silenced-TE transcripts did not involve significantly younger genes than Stable-TE transcripts.

Together, these data demonstrate through a combination of orthogonal approaches that NMD preferentially regulates transcript isoforms that overlap evolutionarily young TE integrations, with relatively younger splice sites and a higher rate of mutational divergence. Furthermore, Tuned-TEs, permitted but downregulated by NMD, may represent an evolutionary intermediate, given intermediate signals of conservation and mutation throughout the analysis.

### NMD safeguards proteomic diversity

To understand the proteomic consequences of TE exonization events regulated by NMD, we compared domain change events when the exonized TEs encoded for protein (Fig. 4h). Non-canonical Tuned-TE and Silenced-TE transcripts were more frequently associated with domain losses than non-canonical Stable-TE transcripts (Fig. 4i, upper panel). Furthermore, stratifying domain changes by the age and conservation of the TE-adjacent splice site showed a trend towards higher rates of lost domains in older and conserved splice site transcripts (Fig. 4i, lower panel).

We then evaluated possible Functionally Distinct protein Isoforms (FDIs) generated by TE exonization events, which were split into four categories of decreasing confidence based on supporting evidence: Very High (VH), High (H), Medium (M), and Low (L). Interestingly, comparing FDI predictions, particularly ones with VH, H, and M support, indicated that Silenced-TE transcripts were the most likely to encode possible FDIs, followed by Tuned-TEs, then followed by Stable-TEs (Fig. 4j). Altogether, these data suggested the degree of NMD regulation correlated with the functional outcome from the isoform generated.

We then examined the predicted FDI events from Tuned-TE and Silenced-TE transcripts (above the ‘M’ threshold) (Fig. S5a-b). We associated FDI loci NMD ratios with overall survival in LUSC. While several loci were significantly associated with worse or better prognosis, the majority showed no significant association (94%, 3,534 out of 3,747 > 0.05 FDR) and, consequently, there was no collective association (Fig. S5c). Therefore, we examined the list of FDIs for exemplary genes involved in central cellular processes, like the maintenance of genomic integrity. Three such FDIs involved SINE-*Alu* elements; isoforms of DEAD box helicase 49 (DDX49), Nei-like DNA glycosylase 3 (NEIL3), and SET Domain Bifurcated Histone Lysine Methyltransferase 1 (SETDB1), known to epigenetically repress TEs^35,36^. The three isoforms were *DDX49-MIRc/AluSp*, *NEIL3-AluSx*, and *SETDB1-AluSx3* (Fig. 5a), and all encoded truncated proteins that retained at least one annotated domain (Fig. 5b, S5d-e). DDX49-MIRc/AluSp retained only a fraction of its helicase domain, but contained its Q motif, known to regulate RNA binding and helicase activity^37^. The protein encoded by *NEIL3-AluSx* retained an annotated glycosylase domain, but lost C-terminal GRF-type Zinc finger domains that mediate localization specificity in ssDNA binding^38,39^. SETDB1-AluSx3 lost its enzymatic SET domains that enable methyltransferase activity, but retained its three Tudor domains which mediate H3K9 binding^40^.

**Figure 5.**
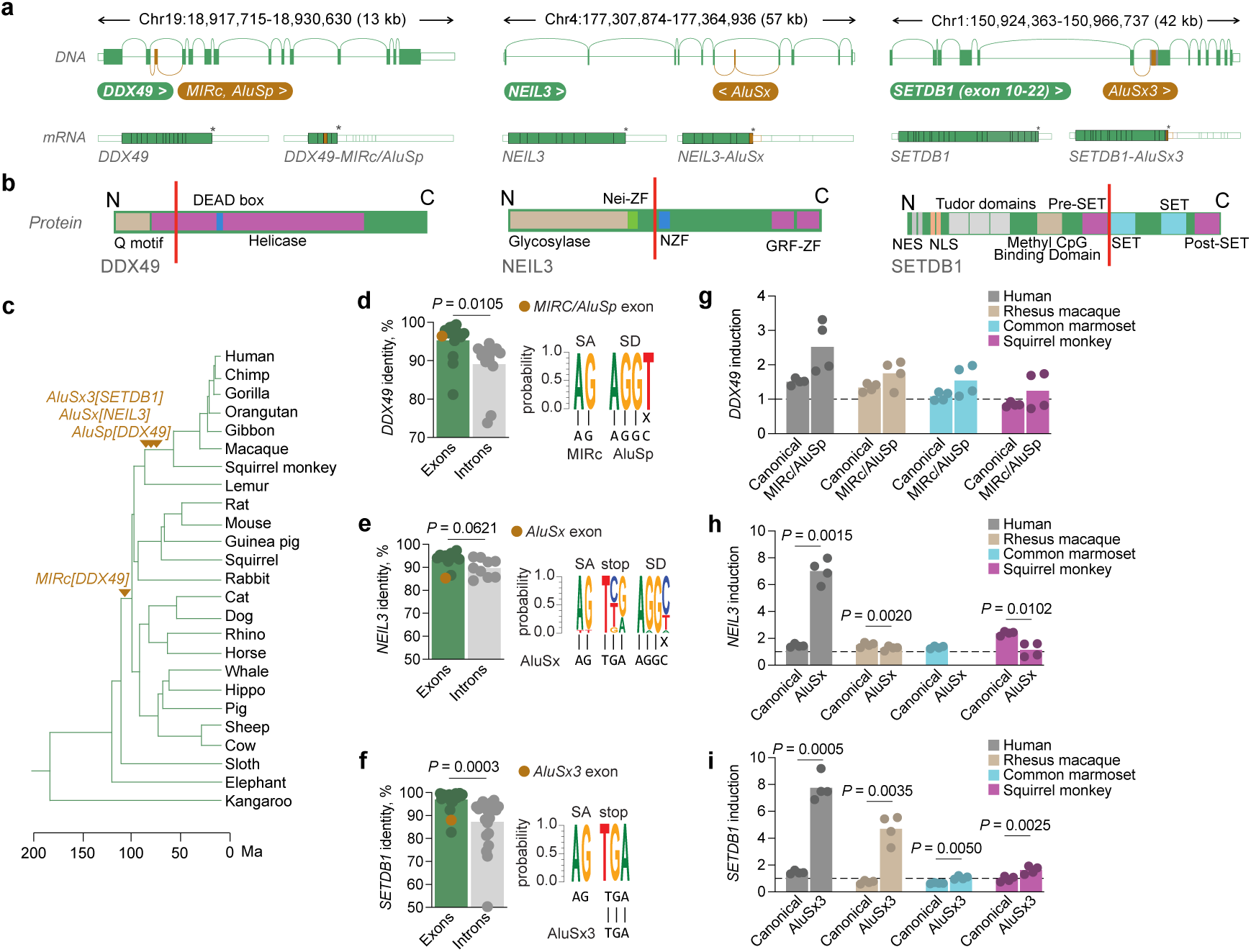
*Alu*-exonizing putative FDI transcripts represent different degrees of evolutionary conservation. **a**, Schematics of the *DDX49*, *NEIL3*, and *SETDB1* (exons 10-22) genes with alternative splicing to intronic *Alu* elements, producing PTC-containing alternative transcript isoforms. **b**, Schematics of the DDX49, NEIL3, and SETDB1 proteins with domains annotated, red vertical lines indicate the approximate truncation position in each respective *Alu* isoform. N, N-terminus; C, C-terminus; ZF, Zinc finger, NZF, Npl4 Zinc Finger; GRF-ZF, Glycine-Arginine-Phenylalanine Zinc finger; NES, Nuclear Export Signal; NLS, Nuclear Localization Signal. **c**, Estimation of the evolutionary age of the TE integrations that generate the *Alu* isoforms based on genomic sequence alignments. **d**-**f**, Mean percent identity of *DDX49* (d) *NEIL3* (e), and *SETDB1* (f) exons and introns across 19 primates, generated by a per-base gap-aware calculation. Symbols represent individual exons and introns, the *Alu* exon is colored differently, and *P* values were calculated using an unpaired two-tailed Student’s *t* test. Adjacent to each bar plot are logo plots for the splice acceptor (SA) and splice donor (SD) sites and stop codon, based on sequence conservation in 19 primate genomes, with the respective sequence in the TE consensus indicated below. **g**-**i**, Induction by RT-qPCR of the canonical and *Alu* isoforms of *DDX49* (g), *NEIL3* (h), and *SETDB1* (i) in Human (HEK293T), Rhesus macaque (LLC-MK2), Common marmoset (CM0203F), and Squirrel Monkey (SQMK-FP) cell lines, relative to HPRT and to DMSO control, after 24 h treatment with 10 µM of eIF4A3-in-1. Symbols represent two technical replicates from two independent experiments. *P* values were calculated with paired two-tailed Student’s *t* tests.

### Evolutionary history of the NMD *Alu* isoforms

All *Alu* elements integrated during and following Happlorhini speciation (Fig. 5c). As a mammal-wide element, the MIRc involved in *DDX49-MIRc/AluSp* was older, although here it aligned only as far back as dogs. Between the three *Alu* isoforms, the MIRc/AluSp exon in *DDX49* was the best conserved. It resembled an exon, and its ORF, putative splice acceptor (SA) and splice donor (SD) sites were perfectly conserved (Fig. 5d, S6a, S6d). The putative SA provided by the *MIRc* element was unchanged from the consensus *MIRc* sequence, but a mutation from the *AluSp* consensus (C to T) provided the putative SD site. Given all primates encoded a ‘T’ at this position, this suggested an immediate mutation in the common ancestor, for instance via an error during the initial integration process.

*NEIL3-AluSx* was the least conserved of the three. The *AluSx* exon resembled an intron (Fig. 5e, S6b, S6f), and only Gorilla, Chimpanzee, and Humans had a mutation from the *Alu* consensus (C to T) that generated a putative SD site. Only humans appeared to encode the PTC (‘TGA’, match to *Alu* consensus) and all other species encoded ‘TCA’ (Fig S6g). Closer inspection of our long read RNAseq data suggested that the ‘TGA’ codon, as annotated in reference hg38 assembly, was not shared by the sequenced cell lines, all of which had a ‘TCA’ codon in this position (Fig S6g), similar to the ancestral allele. Indeed, this PTC appeared to be polymorphic in populations of different genetic ancestry (rs6823747), with higher prevalence (>70%) of the ‘TCA’ codon in individuals of European and African ancestry, but almost exclusive presence of the ‘TGA’ codon in individuals of Asian ancestry (Fig. S6h), suggesting that the acquisition of this PTC is a very recent evolutionary event.

Predicted capacity to produce *SETDB1-AluSx3* was conserved amongst all primates, given perfect conservation of the putative SA site (Fig. 5f, S6b). This SA site did not overlap the *AluSx3* annotation, though species without the *AluSx3* did not have the SA. The nature of the *SETDB1-AluSx3* transcript produced across primates was not well conserved, given a variable ‘TG’ repeat that altered the ORF, generating nevertheless truncated proteins but of different lengths between primates.

We then tested RNA from the HEK293T human cell line, and from three primate cell lines: LLC-MK2 (rhesus macaque), CM0203F (common marmoset), and SQMK-FP (squirrel monkey) for induction of the canonical and *Alu*-exonizing isoforms using the eIF4A3-in-1 inhibitor. As a trend, *DDX49-MIRc/AluSp* was upregulated compared to *DDX49*, but this was not statistically significant (Fig. 5g). *NEIL3-AluSx* was more significantly upregulated compared to *NEIL3* in human cells, in rhesus macaque and squirrel monkey it was significantly less upregulated, and *NEIL3-AluSx* was not detected in common marmoset (Fig. 5h). *SETDB1-AluSx3* was significantly more upregulated compared to *SETDB1* in all species (Fig. 5i).

Altogether, we determined these three NMD-regulated *Alu*-exonisation events represented different levels of evolutionary emergence and conservation. *DDX49-MIRc/AluSp* and *SETDB1-AluSx3* were produced across primates and reasonably conserved, while *NEIL3-AluSx* was much younger and poorly conserved. Conservation status may relate to how reproducibly NMD blockade led to their induction, given *NEIL3-AluSx* was not consistently upregulated compared to *NEIL3*.

### Functional differences in NMD-targeted *Alu* isoforms

All *Alu* isoforms were elevated in HeLa.doubleKD compared to HeLa.scramble, and *DDX49* and *NEIL3* exhibited isoform switching (Fig. S7a). All canonical transcripts were expressed by all lines cancer cell line encyclopedia (CCLE), but *Alu* isoform expression varied (Fig. S7b), and the ratio was in favor of the canonical for all three (Fig. S7c). This was the same for the Normal, LUSC and LUAD data, except for *DDX49-MIRc/AluSp*, which was more highly expressed than canonical *DDX49* in all three cohorts (Fig. S7d). Together these data demonstrated that, despite NMD targeting, all three transcripts were expressed with a capacity to alter host biology dependent on changes in NMD activity.

To determine if the isoforms produced stable proteins, we expressed them as part of a split mNeonGreen system, where the protein of interest was encoded with the 11^th^ chain of mNeonGreen as a C-terminal tag, and transfected alongside a separate construct encoding the 1^st^ to 10^th^ chains, to emit fluorescence upon spontaneous co-localization. All three isoforms and their canonical counterparts emitted mNeonGreen signal detectable by flow and visualized by fluorescence microscopy (Fig. 6a). Next, we overexpressed the isoforms or their canonical counterparts (reported by tdTomato) in HeLa cells and subjected them to short-read RNAseq (Fig. 6b). We checked transfection efficiency by analyzing TdTomato expression (Fig. 6c), which showed equivalent expression of the constructs in each condition, apart from *NEIL3* which was not well expressed.

**Figure 6.**
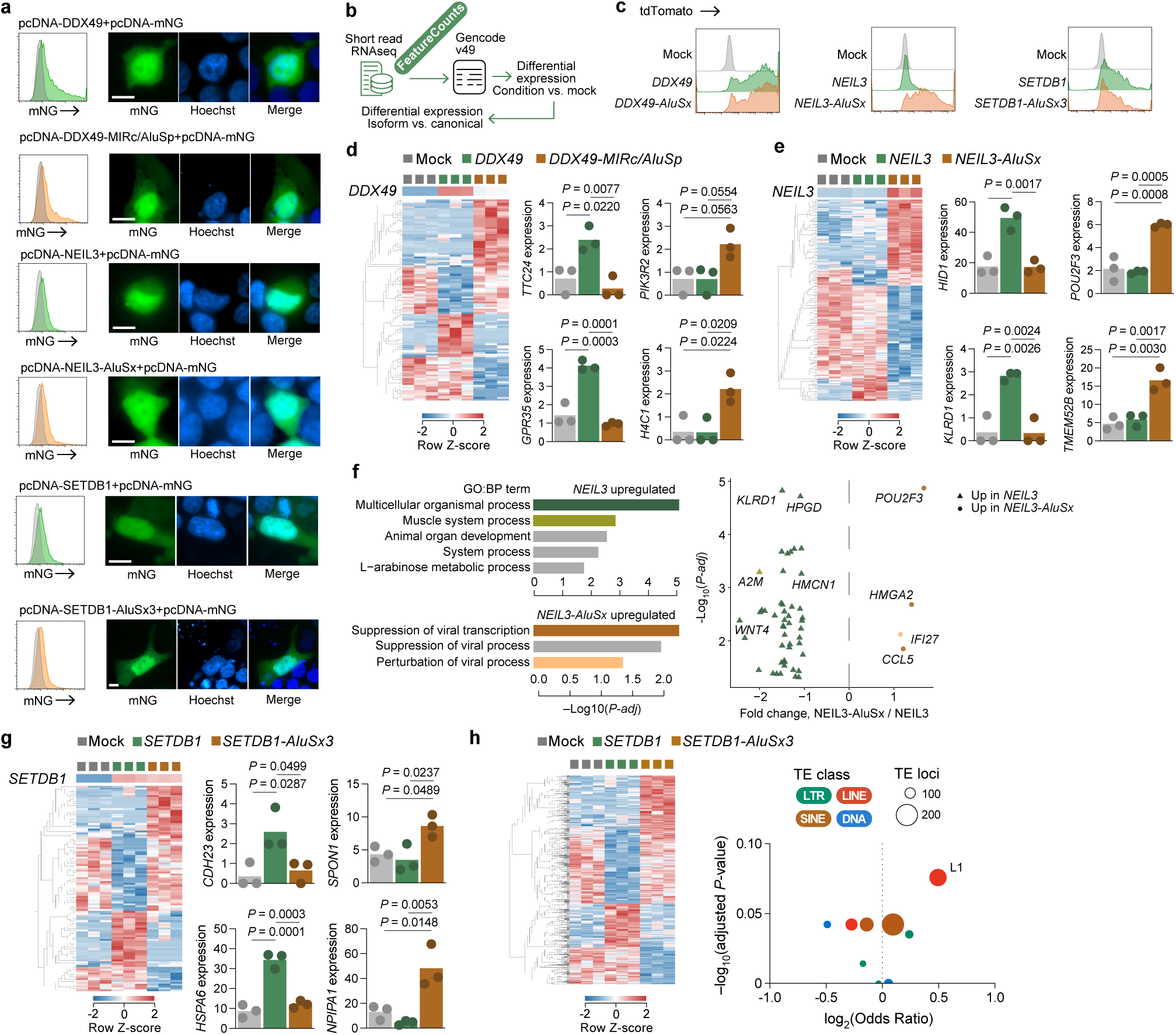
*Alu*-exonizing FDIs encode stable proteins with distinct transcriptional effects. **a**, Flow cytometry (left) and fluorescence imaging (right) of HEK293T cells co-transfected to express canonical and *Alu* isoforms of *DDX49*, *NEIL3*, and *SETDB1* tagged with the 11^th^ chain of mNeonGreen, which associates with the 1^st^-10^th^ chains of mNeonGreen expressed from a different construct (pcDNA-mNG). Hoechst indicates the nuclear stain, and scale bars indicate 10 µm. On the flow cytometry plots, the grey indicates the mock-transfected control, green indicates a canonical transfection, and orange an *Alu* isoform. **b**, Schematic of the analysis approach after short-read sequencing of HeLa cells after transfection with the canonical/*Alu* constructs from (a). **c**, Flow cytometry of tdTomato expression in HeLa cells used for short-read RNAseq after transfection with the canonical/*Alu* constructs from (a) which encode tdTomato from an IRES sequence. **d**, **e**, **g**, Heatmap of transcripts significantly differentially regulated compared to mock, and between each canonical and *Alu* transfection condition (≥2-fold change; *P* ≤ 0.05; *q* ≤ 0.05) after transfection with canonical and *Alu* isoforms of *DDX49* (d), *NEIL3* (e), and *SETDB1* (g), with examples of significantly differentially expressed genes shown on bar plots. Columns represent independent replicates of each condition, and *P* values were calculated with one-way ANOVA. **f**, Left: Top five GO Biological Process terms enriched (g:Profiler, g_SCS-corrected) among genes significantly upregulated in *NEIL3* relative to *NEIL3-AluSx* (top) and in *NEIL3-AluSx* relative to *NEIL3* (bottom). Right: The differentially expressed genes that contribute to a significant GO:BP term in either direction; triangles indicate genes higher in *NEIL3*, circles indicate genes higher in *NEIL3-AluSx*, with exemplary genes labelled. **h**, Left: Heatmap of TEs significantly differentially regulated compared to mock, and between *SETDB1* and *SETDB1-AluS3* transfections. Right: TE family enrichment among significantly upregulated TE loci in *SETDB1-AluS3* compared to *SETDB1* transfection. Each point represents one TE family, point color indicates TE class, and point size indicates the number of TE loci in the given family. Odds ratios and adjusted *P* values were calculated using a two-sided Fisher’s exact test with BH correction, but none were significant.

Then, we performed a differential expression analysis (Fig. 6d-e, g). We confirmed that *NEIL3* was not efficiently expressed after transfection (Fig. 6e), but that all others were detectably elevated (Fig. 6d-e, g). In all cases, there were clusters of genes that were induced selectively by the canonical or *Alu* isoform. *DDX49-MIRc/AluSp* alone upregulated *PIK3R2*, a growth regulator^41,42^, and *H4C1*, a component of the nucleosome^43^ (Fig. 6d).

Despite poor transfection efficiency, *NEIL3* was associated with the unique upregulation of genes, as was *NEIL3-AluSx*, which upregulated *POU2F3*, a transcription factor implicated in small cell lung cancer^44^. Furthermore, *NEIL3-AluSx* was the only isoform to show significant pathway enrichment – it upregulated viral suppression genes (Fig. 6f).

*SETDB1-AluSx3* was upregulated *SPON1*, implicated in lung cancer metastasis^45^, and *NPIPA1*, a nuclear pore complex protein associated with brain development^46^ (Fig. 6g), and with different patterns of TE expression, namely L1 upregulation (although not statistically significant) (Fig. 6h). Given the proposed role for SETDB1 in the repression of L1 elements^47^, this may suggest SETDB1-AluSx3 acted in a dominant negative manner, recruiting endogenous SETDB1 interaction partners away from effective L1 repression.

Altogether, these analyses revealed each isoform had unique functional effects relative to their canonical counterpart proteins, and therefore if expressed, could be subject to evolutionary pressure to stabilize their expression if beneficial, or maintain or enhance their silencing if harmful. In addition, as loci although they were not individually associated with survival outcomes in LUSC, their capacity to modify host biology could complement in the event NMD activity is altered, meaningfully altering the course of disease.

## Discussion

Here, we have determined that NMD acts as a co-translational checkpoint that restrains effects from TE-driven exonisation, complementing the sequence of regulatory barriers known to repress TE activity. Upstream, DNA methylation^22^ and KZFPs^23^ bind TEs and enforce transcriptional silencing, and HUSH-mediated epigenetic repression targets young full length LINE elements^25^ and intronless transcripts^24^. Later in their evolution, TE fragments evolve splice sites^48^ and are subject to splicing repression. *Alu* elements are bound by the RNA-binding protein (RBP) hnRNP C via antisense U-tracts, repressing splicing to over 1,000 cryptic *Alu* exons^26^, most of which were predicted to also introduce frameshifts and PTCs^26,27^. Young LINE elements recruit repressive RBP hubs, and LINE-derived exons that escaped repression were found to be more readily detected upon UPF1 depletion^27^. We show that NMD targeting of TE-exonizing isoforms extends across TE classes and, like the upstream barriers, preferentially acts on the youngest elements.

TE integrations may represent a particularly disruptive, but capacious strategy to drive functional divergence for evolutionary adaptation, given the volume of genetic material introduced through integration events. It may logically follow that NMD is an ideal targeting strategy given its own objective to eliminate complete production of the modified protein product. Here the activity of proteins with outsized effects on their respective pathways is most effectively avoided. Indeed, we have shown that NMD targeted TE-overlapping transcripts are particularly likely, over their stably expressed counterparts, to produce proteins with significant changes in their functional output after the modification of interacting domains. This contrasts with established, NMD-insensitive *Alu* exons, such as in ADARB1, which appear to fine-tune rather than disrupt the activity of the encoded protein^49^.

Perhaps most critical from an evolutionary perspective is that NMD control is incomplete. NMD silenced some TE-overlapping transcripts, but only tuned others, NMD-targeted TE isoforms were detectable at steady state in cell lines, normal lung and lung tumors, and the three *Alu* isoforms we examined produced stable proteins with distinct functional effects. Independently, translated peptides and ribosome profiling evidence were found for a subset of putative NMD-inducing *Alu* exons^49^, and inclusion of newly derepressed *Alu* exons correlated with downregulation of the host transcripts, consistent with NMD^26^. Our data suggest that this graded control correlates with an evolutionary trajectory: Silenced-TEs were the youngest, most diverged and least conserved, Stable-TEs the oldest and most conserved, and Tuned-TEs intermediate throughout. We propose that TE-derived exons pass through layered checkpoints: splicing repression determines whether TE sequence enters mature mRNA, NMD then titrates how much of a PTC-containing isoform is translated, and selection acts on mutations that shift NMD-susceptibility. Only isoforms that escape both layers become established coding exons^49^. This extends the model in which hnRNP C repression allows Alu elements to evolve near-neutrally until leaky exonization exposes them to selection^26^. NMD may therefore extend the period over which new TE-derived protein sequence can be tested by selection, beyond the point at which splicing-level repression has been relieved.

It is challenging to qualify the physiological effects of changes to a process as broad as NMD. The role of NMD in cancer in particular may be dual and context dependent. Tumors exploit NMD to silence transcripts carrying truncating mutations in tumor suppressor genes and the efficiency of NMD shapes which nonsense mutations are selected during tumor evolution^9^. NMD components are also recurrently altered in some tumors, most notably through somatic UPF1 mutations in pancreatic adenosquamous carcinoma^50^. In addition, NMD activity can be suppressed by stresses in the tumor microenvironment, such as hypoxia and nutrient deprivation acting through eIF2α phosphorylation, which can promote tumor growth^51,52^.

Direct studies of NMD in lung cancer remain limited, although reduced UPF1 expression in lung adenocarcinoma has been linked to activation of TGF-β signaling and epithelial-mesenchymal transition^53^. NMD further shapes tumor immunogenicity. Frameshift-derived transcripts that escape NMD are associated with response to immune checkpoint blockade^11,54^, and targeted NMD inhibition in tumor cells induces non-canonical antigen expression and tumor immunity in mice and patient-derived models^10,11^.

We observed that NMD activity is increased in lung tumors, together with the upregulation of NMD pathway proteins, and lower activity associated with poorer survival specifically in LUSC. This suggests that the consequences of NMD activity differ between histological subtypes. TE-derived NMD substrates, which previous analyses have not considered, may contribute to these differences, as well as to the creation of non-canonical tumor-associated antigens, further supporting the notion of selective NMD inhibition as a potential therapeutic intervention^54^.

More broadly, our study reveals a significant and unexplored reservoir of TE-driven transcript isoforms that is tamed and evolved by NMD. Time and again, we and others have demonstrated how the functional pathway integration of such isoforms changes biological outcomes with meaningful consequences. A better understanding of this overlooked and evolving transcriptome will be essential to more comprehensively understand cellular behavior in health and disease.

## Methods

### Cell culture & transfections

HEK 293T (RRID:CVCL_0063), A-549 (RRID:CVCL_0023), HCC4006 (RRID:CVCL_1269), NCI-H441 (RRID:CVCL_1561), Calu-1 (RRID:CVCL_0608), HT-1080 (RRID:CVCL_0317), HeLa (RRID:CVCL_0030), SQMK-FP (RRID:CVCL_6451), LLC-MK2 (RRID:CVCL_3009), and CM0203F^55^ were obtained from the Cell Sciences facility of the Francis Crick Institute, verified mycoplasma free, and validated by DNA fingerprinting (human lines only). Cells were cultured in Dulbecco’s modified Eagle’s medium, Roswell Park Memorial Institute 1640 medium, or Iscove’s modified Dulbecco’s medium, supplemented with L-glutamine (2 mM) (Thermo Fisher Scientific), penicillin-streptomycin (50 U/ml) (Gibco, cat # 15070063), and 5 or 10% heat-inactivated fetal bovine serum (FBS) (Gibco), as indicated in the table below. The eIF4A3-in-1 compound (Tebubio, cat # T11170) was dissolved in DMSO prior to treatment at 10 µM for 24 h. The spliced coding sequences for the canonical and *Alu* isoforms of *DDX49*, *NEIL3*, and *SETDB1* were synthesized with a C-terminal mNeonGreen tag (11^th^ chain) and inserted into the pcDNA-IRES-tdTomato vector. For co-localization assays, the 1^st^ to 10^th^ chains of mNeonGreen were synthesized and inserted into the pcDNA vector for co-transfection. Transfections were performed for 48 h, using GeneJuice transfection reagent (mock was GeneJuice only) (Sigma-Aldrich, cat # 70967). GenScript performed gene synthesis and cloning, and construct sequences were validated by sequencing via Plasmidsaurus.

| Cell line | Species | Cell Origin | Media |
| --- | --- | --- | --- |
| HEK293T | Human | Embryonic kidney | IMDM + 5% FBS |
| HeLa | Human | Cervical adenocarcinoma | DMEM + 10% FBS |
| A-549 | Human | Lung adenocarcinoma | DMEM + 10% FBS |
| HCC4006 | Human | Lung adenocarcinoma | RPMI + 10% FBS |
| NCI-H441 | Human | Lung adenocarcinoma | RPMI + 10% FBS |
| Calu-1 | Human | Lung squamous cell carcinoma | DMEM + 10% FBS |
| HT-1080 | Human | Fibrosarcoma | DMEM + 10% FBS |
| SQMK-FP | Squirrel monkey | Epithelial kidney | DMEM + 10% FBS |
| LLC-MK2 | Rhesus monkey | Epithelial kidney | DMEM + 10% FBS |
| CM0203F | Common marmoset | Bone marrow stromal cell | DMEM + 10% FBS |

### Flow cytometry

Cell suspensions were analyzed on a BD Biosciences LSRFortessa using FACSDiva v8.0, and analyzed using FlowJo v10. Gating strategies are provided in Fig. S8. Cells were washed with and resuspended in 1% (w/v) bovine serum albumin (BSA) in phosphate-buffered saline (PBS) for acquisition.

### Fluorescence imaging

Before imaging, cells were grown on 1.5 glass slips (VWR, cat # 631-0153), then fixed with 10% neutrally buffered formalin for 15 min on ice. Samples were then stained with 5 µg/mL Hoechst 33342 (Thermo Fisher Scientific, cat # 62249), before being washed in 1% (w/v) BSA in PBS. Sample coverslips were then sealed to SuperFrost glass slides (Hawksley, cat # FD-MS-SUPER) using Vectashield (2B scientific, cat # H-1000-10), and imaged on a ZEISS Apotome 3 using ZEN software.

### RNA preparation, PCR, and in-house sequencing

RNA was extracted from cell lines using the RNeasy kit with DNAse treatment according to the manufacturer’s protocol (Qiagen, cat # 74104). Short-read library prep was performed with the NEBNext Ultra II Directional PolyA mRNA kit (New England Biolabs, cat # E7760) before sequencing on a NovaSeq (Illumina). For long-read sequencing, cDNA libraries were prepared using the Oxford Nanopore Technologies (ONT) PCR-cDNA Sequencing Kit V14 (SQK-PCB114.24) according to the manufacturer’s protocol, with 500 ng of total RNA input per sample and 20 PCR amplification cycles. Equimolar or rebalanced sample pools were prepared using Qubit fluorometric quantification and TapeStation D5000 analysis, ligated with ONT Rapid Adapters, and sequenced across PromethION flow cells on a P2 Solo device. Basecalling was performed on a HPC cluster using Dorado (v1.1.1) with model dna_r10.4.1_e8.2_400bps_sup@v5.2.0. For RT-qPCR, extracted RNA was synthesized into cDNA with the High-Capacity Reverse Transcription Kit (Thermo Fisher Scientific, cat # 4368813), before qPCR using Fast SYBR Green master mix (Thermo Fisher Scientific, cat #EP0701) on a QuantStudio 3 (Applied Biosystems). Gene-, isoform-, and species-specific primers (table below) were designed using primer3plus. It should be noted differences in amplification efficiency of primate cDNA with the human *HPRT1* primer pair meant inter-species transcript expression was not comparable. The induction values in Fig. 5g-i were calculated by dividing the technical replicate 2^−ΔCt^ values (eIF4A3-in-1/DMSO).

### PCR primers

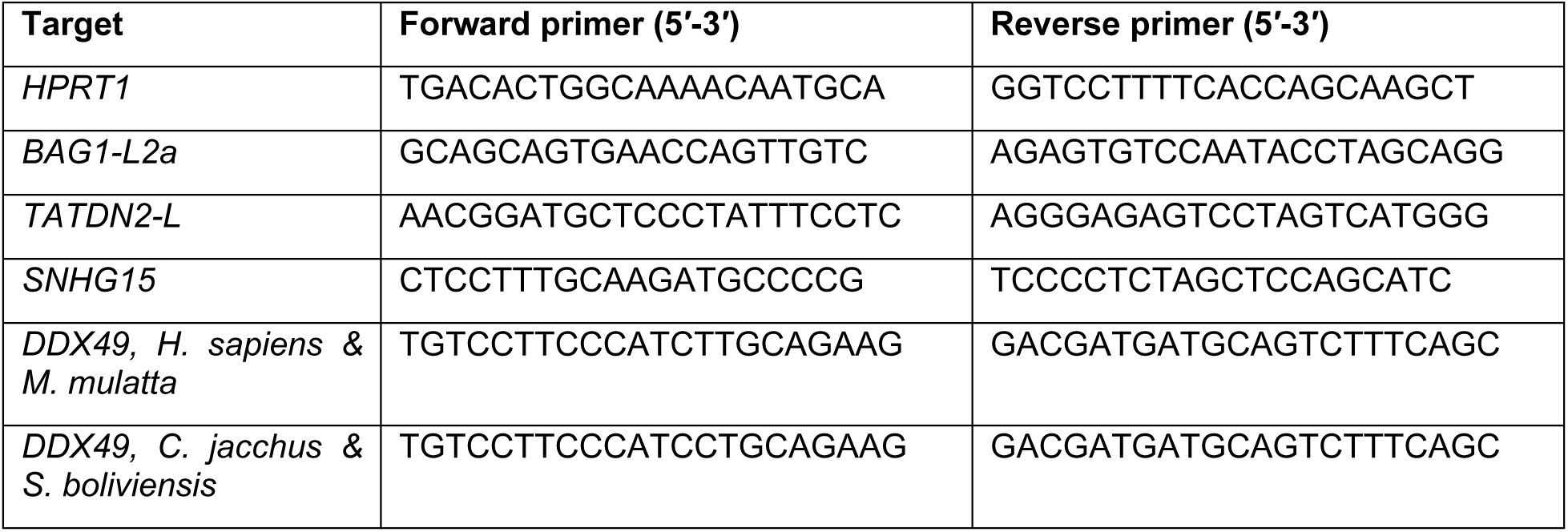
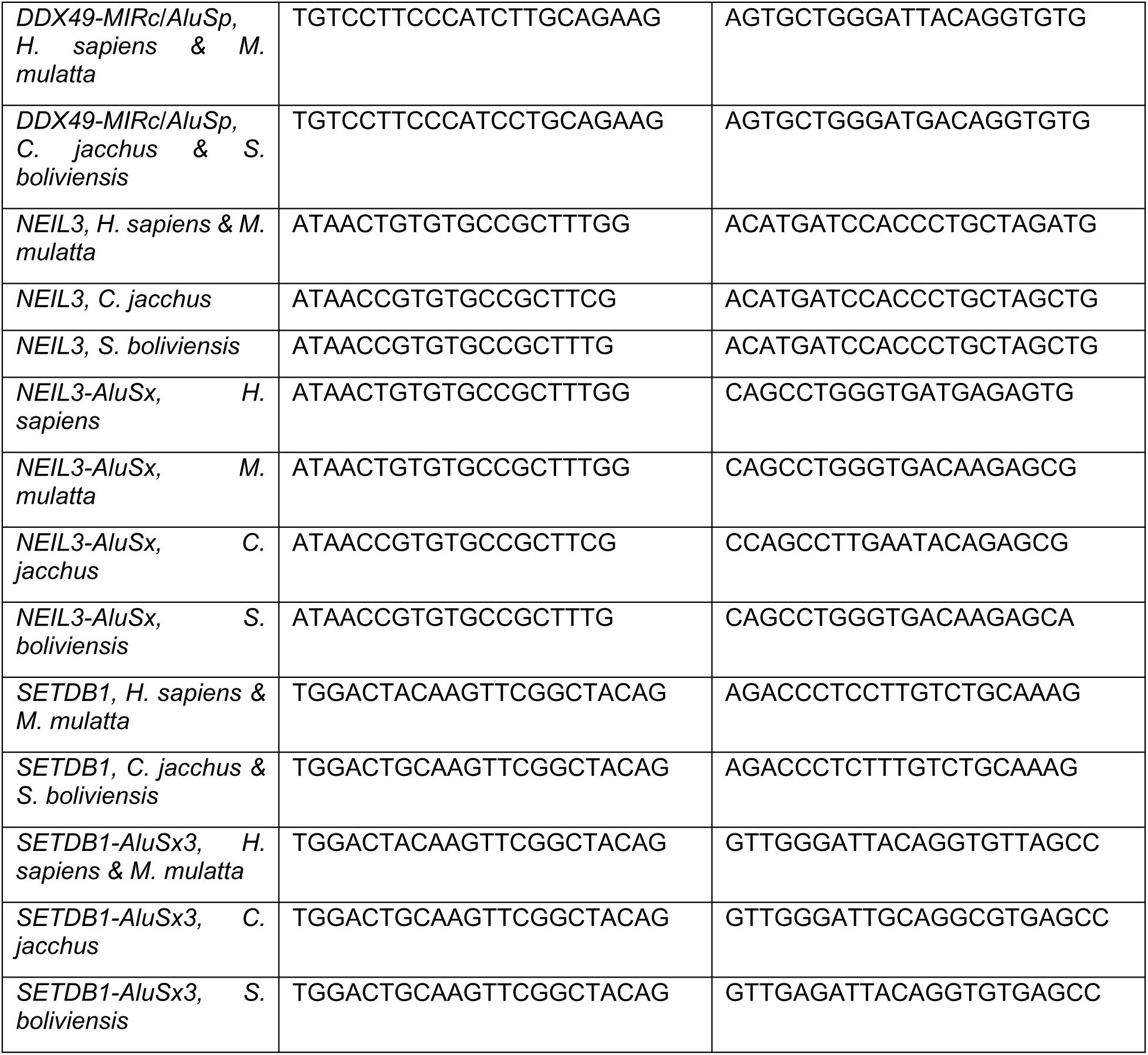

### Transcriptomic assembly

Raw ONT reads were mapped to hg38 using minimap2 v2.26 in long read mode (-ax splice), generating per sample .bam files. These were used to generate per sample .gtf transcriptome files using StringTie v3.0.1 in ‘max capture’ mode (coverage 0.01, isoform fraction 0, junction coverage 1, no trimming, minimum anchor length of 8, long read mode, and with a clean and collapse read approach, no assembly). Processed per sample .gtf files were annotated using Gffcompare v0.12.10, with built-in class codes used for splice pattern analyses. Comparing the transcriptomes on a per sample basis was done with random downsampling to account for different read depth.

Per sample .gtf files were collapsed and assembled using StringTie in merge mode (coverage 0, FPKM/TPM threshold 0), with variable isoform fraction cutoff settings (0.01% to 10%). Varying this threshold, we attempted to quantify the balance between over-assembly and over-stringency using a panel of parameters, including the total number of transcripts assembled, the average number of isoforms per locus, and a variability coefficient which measured the evenness of read count distribution across each locus’s isoforms. However, all these scaled in a near-linear fashion, and so did not discriminate an optimal threshold. In contrast, comparing the proportion cutoffs by how many genes appeared in the final assembly with over 20 isoforms, a signal of per gene diversity, showed an exponential drop-off from ∼6k genes at a 0.1% proportion cutoff to ∼1.5k genes at a 1% proportion cut-off. Therefore, we used a 1% proportion cutoff to generate the final assembly. Our NET .gtf file was also annotated using Gffcompare and annotated on a per exon and per transcript basis for repeat overlap using a custom pipeline. If not generated in-house, RNAseq reads were obtained from the TCGA^56^, CCLE^57^ or the GEO repository.

### TE annotation and processing

We annotated TEs in hg38 using RepeatMasker v4.2.3 and the Dfam3.9 consensus library. We identified potential sources of overestimation in this raw annotation, that could later manifest in the TE-transcript overlap counting. These occurred due to low scoring gaps breaking up integrations or being assigned as closely related but different TE families, and because ERVs appear separately as internal and LTR sequences in the Dfam3.9 library. In an extreme case, a single HERVK14_C4 integration appeared as ten separate annotations (Fig. S3b).

To remedy this, we performed a set of Python v3.6.8 computations. Firstly, a ‘rename competing annotations’ step, which found adjacent members of the same TE family (e.g. ERVK for HERVK and HERVK14_C4), and renamed them based on the longest and highest scoring annotation for the region (RepeatMasker Smith-Waterman scoring). Next, we used a ‘merge adjacent fragments’ step to join neighboring annotations of identical repeat subfamilies when separated by a genomic gap less than 50 bp. This step not only combined subfamily annotations generated by the ‘rename competing annotations’ step, but also any adjacent subfamily annotations that were fragmented as a result of low consensus match scoring over a short region. In these cases, RepeatMasker stringency settings had split what was likely one initial integration into two or more separate elements with unannotated gaps, increasing the number of total annotations. Finally, ERV annotations were collapsed, merging neighboring LTR and internal regions based on a custom lookup table (table S1), determined by parsing the curated Dfam3.9 annotation file.

### Read alignment and quantification

For quantification, reads were aligned and quantified against our NET assembly or Gencode v49 using Salmon v1.10.0, after generating a Salmon-compatible transcript index (TCGA, CCLE and public HeLa/HT1080 datasets), or aligned using HISAT2 v2.2.1 and quantified using featureCounts v1.6.4 against Gencode v49 or the processed Dfam3.9 annotation (overexpression experiments). TPM matrix files or counts tables from this step were analyzed either in Qlucore (t-SNE, heatmap, principal component analysis with ranked variance testing, two-group comparisons), or in R with DESeq2 for differential expression analysis, followed by pathway analysis using g:Profiler^58^. For the heatmaps in Fig. 6, differentially expressed genes were identified using Qlucore, by first comparing each overexpression condition relative to Mock (two-group t-test, *P* < 0.05, fold change ≥ 2), then comparing each canonical protein with the respective *Alu* isoform within the union of these genes using the same thresholds.

For the TE analysis in Fig. 6h, multi-mapping reads were assigned to a single locus, and TE counts were normalized using DESeq2 size factors. Differentially expressed TE loci between *SETDB1-AluSx3* and *SETDB1* were identified in Qlucore (two-group t-test, P < 0.05, fold change ≥ 2), restricted to loci altered by either construct relative to Mock.

### ORFs and NMD prediction

TranSuite predicted primary ORFs in transcripts from the NET assembly based on any overlapping start codon with an annotated counterpart transcript, or by prioritizing the longest ORF. NMD-targeting was then predicted using TranSuite’s TransFeat module; by identifying the primary ORF’s stop codon and whether it sat at least 50 nucleotides upstream of the last exon-exon junction complex, if there were any alternate or upstream ORFs, or a long 3’ UTR. To assess the distribution of NRTs, unique SINE, LINE, LTR and DNA loci overlapping NMD-TEs or Stable-TE transcripts (chr1 to 22, and X) were assigned by midpoint to non-overlapping 1 Mb windows.

### Per donor NMD ratio, and NMD composite expression score calculations

Per locus NMD scores were calculated by dividing the expression of NRTs by total expression from all transcripts from that locus (6,408 loci total), which were then averaged per donor.

To generate the NMD composite expression score, we summed the TPM for all NET assembly transcripts from *UPF1*, *UPF2*, *SMG1*, *SMG5*, *SMG6*, *SMG7* and *eIF4A3*, filtering out any transcripts expressed below 0.1 TPM. Expression values were then log_10_-transformed (TPM + 0.01) and z-scored per gene across all donors from the Normal lung, LUAD, and LUSC cohorts. Per donor composite scores represented the mean of the seven gene z-scores.

### Evolutionary analyses

TE integration ages were estimated by lifting TE coordinates from hg38 to ten species using UCSC chain files (chimpanzee, gorilla, orangutan, gibbon, rhesus macaque, marmoset, mouse lemur, tree shrew, mouse and dog), assigning each TE to the age interval between the most distant species to which it lifted and the next. For each transcript, the youngest overlapping TE was used. Sequence divergence (milliDiv) was obtained from RepeatMasker output. Splice sites were dated by lifting the donor and acceptor dinucleotides of each intron to eight species and checking that the canonical GT/AG sequence was retained in the target genome. Each site was assigned the divergence time of the most distant species in which it was preserved, and conservation was scored with phastCons100way. For the joint age-conservation analysis, sites were classified as old or young (≥43 or <43 Ma) and as conserved or non-conserved (mean phastCons ≥0.5 or <0.5). Gene ages were assigned by phylostratigraphy using phylomapr.

### Sequence alignments, splice motif identification, and comparative genomics

Genomic sequence alignments were obtained from Ensembl^59^ for the *Alu* isoform comparisons for the following 19 primate species: *Homo sapiens*, *Pan paniscus*, *Pan troglodytes*, *Gorilla gorilla*, *Pongo abelii*, *Nomascus leucogenys*, *Cercocebus atys*, *Mandrillus leucophaeus*, *Papio anubis*, *Macaca fascicularis*, *Macaca mulatta*, *Macaca nemestrina*, *Chlorocebus sabaeus*, *Rhinopithecus bieti*, *Rhinopithecus roxellana*, *Aotus nancymaae*, *Callithrix jacchus*, *Cebus imitator*, and *Saimiri boliviensis boliviensis*. Splice site motifs were identified and visualised using the WebLogo tool^60^. For liftOver analyses, chain files were obtained from the University of California Santa Cruz resource^61^ for the following species: *Pan troglodytes* (PanTro6), *Gorilla gorilla* (GorGor6), *Pongo abelii* (PonAbe3), *Nomascus leucogenys* (NomLeu3), *Macaca mulatta* (RheMac10), *Callithrix jacchus* (CalJac4), *Microcebus murinus* (MicMur3), *Tupaia chinensis* (TupChi1), *Mus musculus* (Mm39), and *Canis lupus familiaris* (CanFam6). For each region and species, gap-aware identity was calculated as the proportion of human bases that were both aligned and matched in the other species: 100 × (1 − (mismatches + species gaps) / human bases). Intron/exon-species pairs that were over 75% gapped while an adjacent intron/exon aligned below 25% gaps were treated as likely alignment artefacts and excluded. Identity was then averaged across species per exon or intron.

### Protein structure prediction and visualization

Protein sequences were obtained from UniProt^62^ or translated using SnapGene^63^, predicted by AlphaFold^64^, and visualized using the Research Collaboratory for Structural Bioinformatics Protein Data Bank Mol* 3D viewer^65^.

### Functionally Distinct Isoform predictions

Firstly, ORFs of non-Gencode Stable-TE, Tuned-TE and Silenced-TE transcripts (retrieved from the TranSuite prediction) were translated and Ensembl canonical counterpart proteins were retrieved. Then, we compared protein domains in the primary sequence of the Stable-TE/Tuned-TE/Silenced-TE proteins, which were identified via sequence homologs using the HMMER algorithm and the Pfam domain library. In this context, Stable-TE transcripts were filtered for those that were unannotated in Gencode–this group therefore represented previously undescribed ‘evolutionarily tolerated’ TE exonization events that modified the canonical protein. FDI events were predicted by passing the HMMER-generated protein domain predictions through the 3did^66^ and DOMINE^67,68^ interaction databases. FDI candidates were predicted based on retention of an interacting domain combined with loss of an interacting domain, and Very High confidence evidence was assigned to interactions derived from 3did, as the database contained crystal-structure-based evidence. High, medium-confidence, and low-confidence interactions were derived from the intrinsic computational prediction values assigned in the DOMINE database.

### Software packages for analysis

Analysis was performed in Rstudio v2025.05.1+513 running R v4.5.1. Packages used were DESeq2, data.table, dplyr, GenomicRanges, Biostrings, rtracklayer, biomaRt, phastCons100way.UCSC.hg38, GenomicScores, gProfiler, phylomapr, circlize, httr, jsonlite, survival, survminer and eulerr. Resources used were HMMER v3.4, Pfam, 3did, and DOMINE.

### Statistical analyses

Statistical comparisons were performed in GraphPad Prism 10 or in R. When data was normally distributed, groups were compared using paired or unpaired two-tailed Student’s *t* tests or using one-way ANOVA. For non-normally distributed data, groups were compared using two-sided Wilcoxon rank-sum (Mann-Whitney U) tests, or Kruskal-Wallis tests with Dunn’s test for post-hoc comparisons, with multiple-testing correction. Benjamini-Hochberg (BH) correction was applied for tests performed across categories. For categorical data, groups were compared using chi-square tests, and enrichment was measured using Fisher’s exact tests with BH correction. Survival association was tested using Kaplan-Meier analysis, using the log-rank Mantel-Cox test, and for multivariate analysis, Cox proportional-hazards regression testing. The optimal-cutpoint was a Cox (Wald) survival analysis with false-discovery rate correction. Gene Ontology Biological Process term enrichment was performed using g:Profiler with g:SCS multiple-testing correction.

## Acknowledgements

We thank Callum Goodale-Hall, Jean-David Larouche and other members of the Retroviral Immunology lab for their helpful scientific discussion and support. We are grateful for assistance from the Advanced Sequencing, Cell Services, Flow cytometry, Advanced Light Microscopy and Scientific Computing facilities at the Francis Crick Institute.

## Data availability

The RNAseq data generated as part of this study are deposited with the EMBL-EBI repository (E-MTAB-17773, E-MTAB-17775). Other public short-read RNAseq data used in this study were from the LUSC and LUAD cohorts of the TCGA^56^, CCLE^57^, SMG7 KO HT-1080 cells (GSE232333)^32^, and HeLa cells transfected with KD siRNA against UPF1, SMG6, SMG7 or combined SMG6 and SMG7 (GSE86148)^29^ and long-read RNAseq data from HeLa cells transfected with KD siRNA against UPF1, SMG6, SMG7 or combined SMG6 and SMG7 (E-MTAB-10452)^28^.

