## Supplemental_figures for "Nonsense-mediated decay constrains transposable element-driven evolutionary innovation"

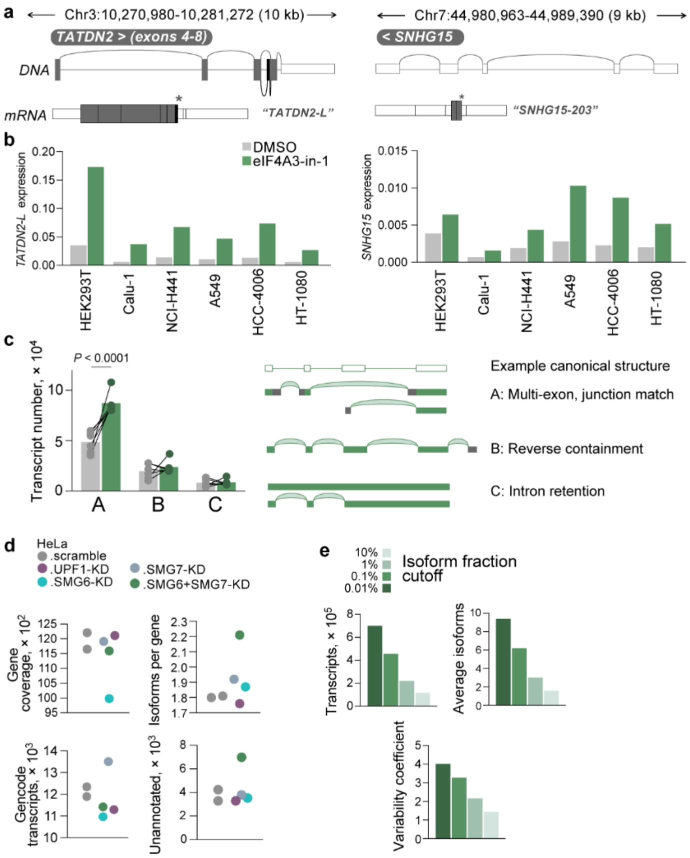


**Figure S1 | eIF4A3 inhibition phenocopies double SMG6/SMG7 knockdown by increasing isoform diversity. a**, Schematic of the *TATDN2* and *SNHG15* genes, with alternative splicing in *TATDN2* to produce the PTC-containing long transcript isoform (*TATDN2-L*), and a putative PTC-containing ORF in *SNHG15*. **b**, RT-qPCR of *TATDN2-L* and *SNHG15* expression in cell lines treated with DMSO or 10 μM of eIF4A3-in-1 inhibitor for 24 h, relative to HPRT. **c**, Number of transcripts in the DMSO or eIF4A3-in-1-treated cell lines per structural category. The *P* value was calculated with a two-tailed paired Student's *t* test. **d**, Gene coverage, isoforms per gene, Gencode, and unannotated transcripts in long-read HeLa RNAseq data[28]. **e**, Transcript number, average isoforms, and variability coefficient of versions of NET assembly, varying the isoform fraction cutoff threshold.

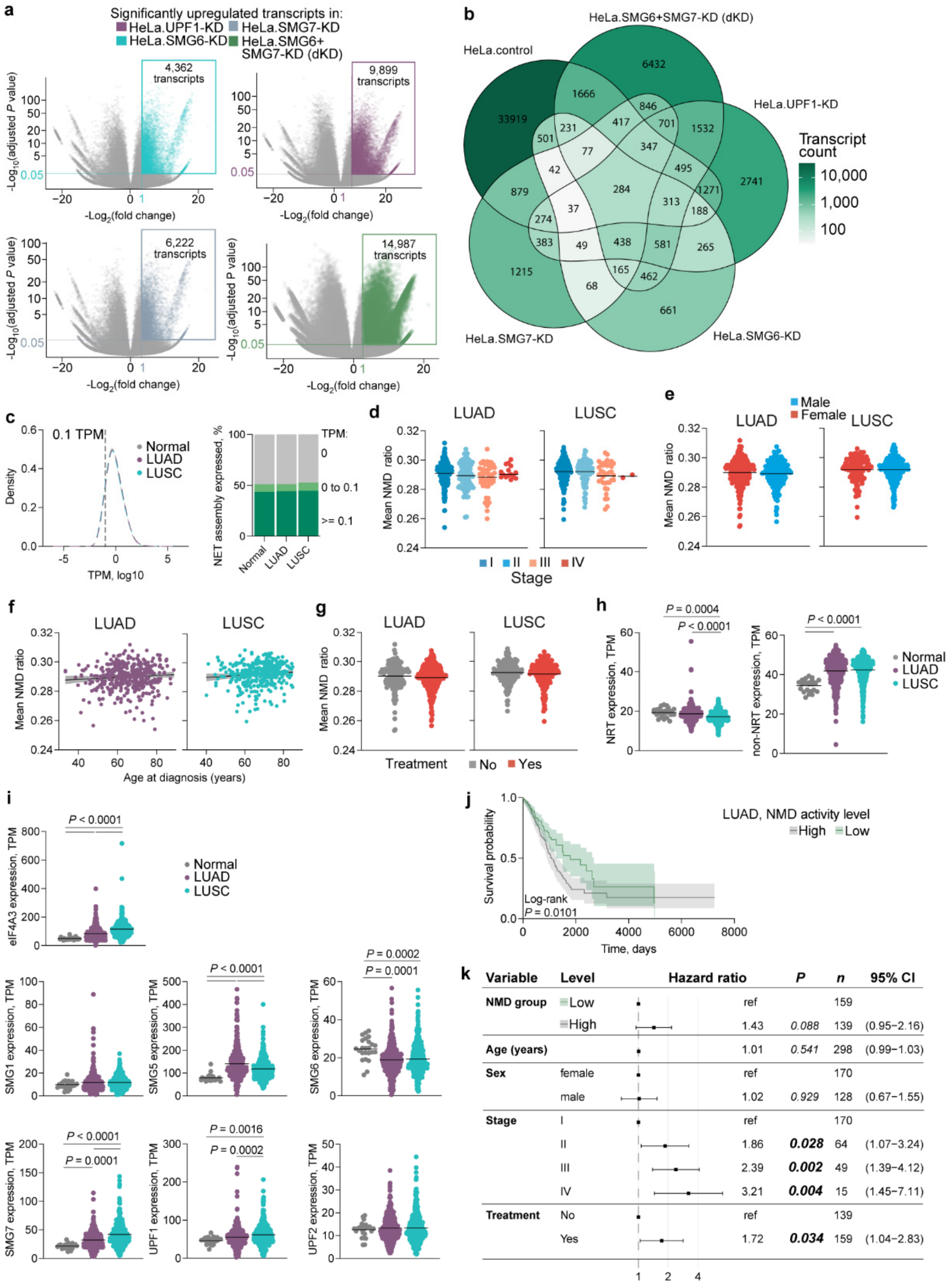


**Figure S2 | NMD inhibition has mostly overlapping effects, and its activity does not depend on lung cancer patient variates. a**, Significantly upregulated transcripts (≥2-fold change; $P \leq 0.05$; $P.\ adj \leq 0.05$) after KD with NMD pathway proteins compared to HeLa.control (transcripts above 1 TPM in HeLa.scramble). **b**, Venn diagram overlap (with numbers) of the transcripts upregulated by each KD condition and compared to HeLa.control. **c**, Left: Expression distribution in TPM of transcripts detected in Normal lung, LUAD, and LUSC TCGA cohorts, with dashed line indicating 0.1 TPM cutoff used for further analyses. Right: Proportion of NET assembly expressed by Normal lung, LUAD, and LUSC cohorts. **d-g**, Mean NMD ratio in LUAD and LUSC cohorts stratified by tumor stage (d), sex (e), age at diagnosis (f), treatment status (g). Symbols represent individual donors. **h**, **i**, NRT (h, left), non-NRT (h, right) and core NMD pathway protein (i) expression in the same cohorts as (c). Symbols represent individual donors, and *P* values were calculated using a Kruskal-Wallis test followed by Dunn's test for pairwise post-hoc comparisons with multiple-testing correction. **j**, Kaplan-Meier plot of overall survival probability in LUAD patients stratified into two groups by median NMD activity level. The *P* value was calculated with a log-rank Mantel-Cox test. **k**, Forest plot of a multivariate Cox proportional hazards model for overall survival in LUAD patients, with NMD activity group (low vs. high), age, sex, tumor stage (I-IV), and treatment status included as covariates. Squares indicate hazard ratios and error bars indicate 95% confidence intervals relative to each variable's reference level. *n* indicates the number of patients in each group.

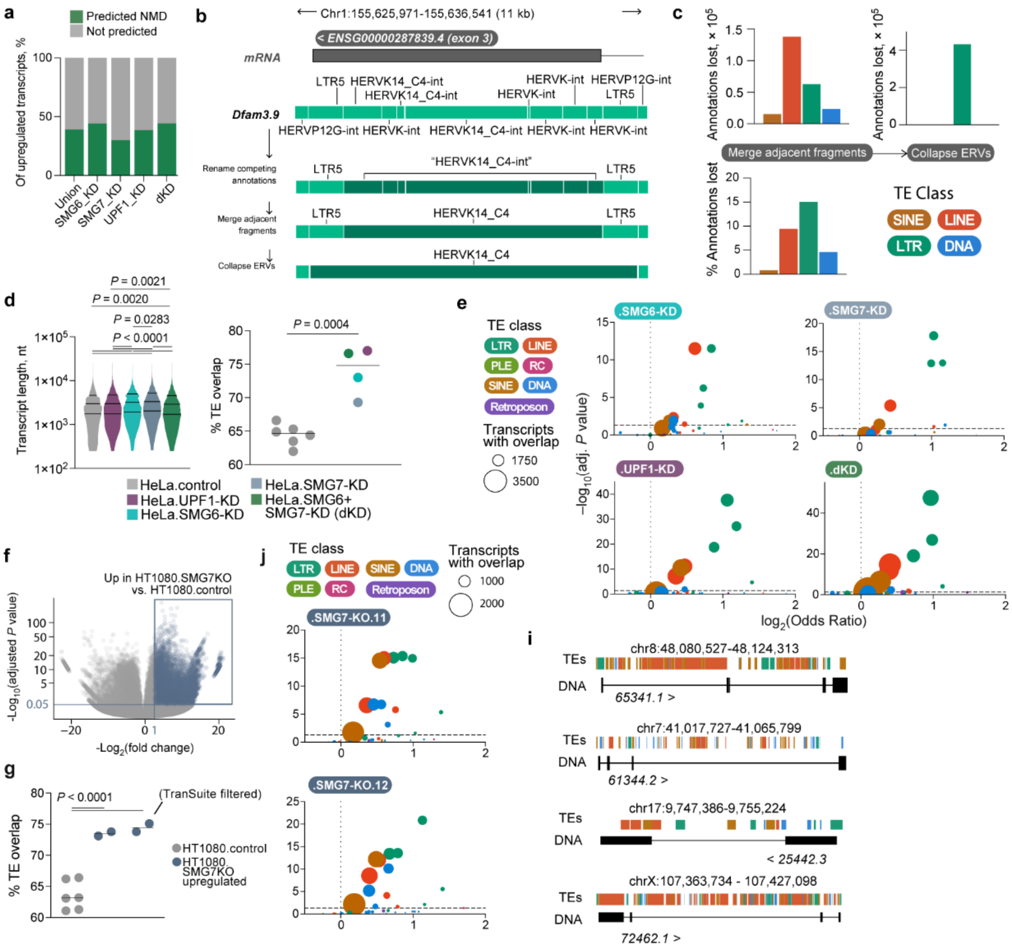

**Figure S3 | NMD has a significant and reproducible tendency to regulate TEs, in particular LTR elements. a**, Proportion of transcripts significantly upregulated by each of the NMD KD conditions that were predicted to be NMD-targeted by TranSuite. **b**, Schematic in a HERVK14_C4 locus illustrating the annotation fragmentation issue across RepeatMasker Dfam3.9, and how we sequentially remedied this. **c**, Above: Total TE annotations removed by class through two pipeline steps. Below: The proportion of per TE class annotations lost. **d**, Left: Length of the transcripts upregulated by each NMD KD. *P* values were calculated using a Kruskal-Wallis test followed by Dunn's test for pairwise post-hoc comparisons with multiple-testing correction. Right: Mean proportion of transcripts overlapping a TE in HeLa.control and those significantly upregulated by each NMD KD when selecting for transcripts of matching length. Symbols represent individual replicates (HeLa.control) or each different NMD KD. *P* values were calculated with a two-tailed unpaired Student's *t* test. **e**, TE family enrichment among TEs overlapping transcripts upregulated by each NMD KD condition, relative to TE-overlapping transcripts in HeLa.control. Each point represents one TE family, point color indicates TE class, and point size indicates the number of transcripts with that TE family overlap. Odds ratios and adjusted *P* values were calculated using a two-sided Fisher's exact test with BH correction. The dashed horizontal line marks adjusted *P* = 0.05, and the dashed vertical line marks an odds ratio of 1. **f**, Significantly upregulated transcripts (≥2-fold change; $P \leq 0.05$; $Padj \leq 0.05$) in HT1080.SMG7KO cells compared to HT1080.WT. **g**, Mean TE overlap proportion in HT1080.SMG7KO cells and HT1080.WT, and filtering for those predicted to be NMD targeted by TranSuite. Symbols represent individual replicates, and *P*

values were calculated with one-way ANOVA. **h**, TE family enrichment among TEs overlapping transcripts upregulated in HT1080.SMG7KO, relative to TE-overlapping transcripts in HT1080.WT. Each point represents one TE family, point color indicates TE class, and point size indicates the number of transcripts with that TE family overlap. Odds ratios and adjusted *P* values were calculated using a two-sided Fisher's exact test with BH correction. The dashed horizontal line marks adjusted $P = 0.05$, and the dashed vertical line marks an odds ratio of 1. **i**, Previously undescribed transcripts from unannotated genomic regions, including transcript 65341.1, a five-exon mRNA from chromosome eight driven by a MER21B LTR element, and 61344.2, a four-exon mRNA from chromosome seven derived in part from an MLT1L LTR element. As well as transcript 25442.3, a two-exon mRNA from chromosome 17 initiated by an MLT1B LTR element and exonizing two downstream L1M elements, and transcript 72462.1, a four-exon mRNA from the X chromosome driven by a L1PA2 element.

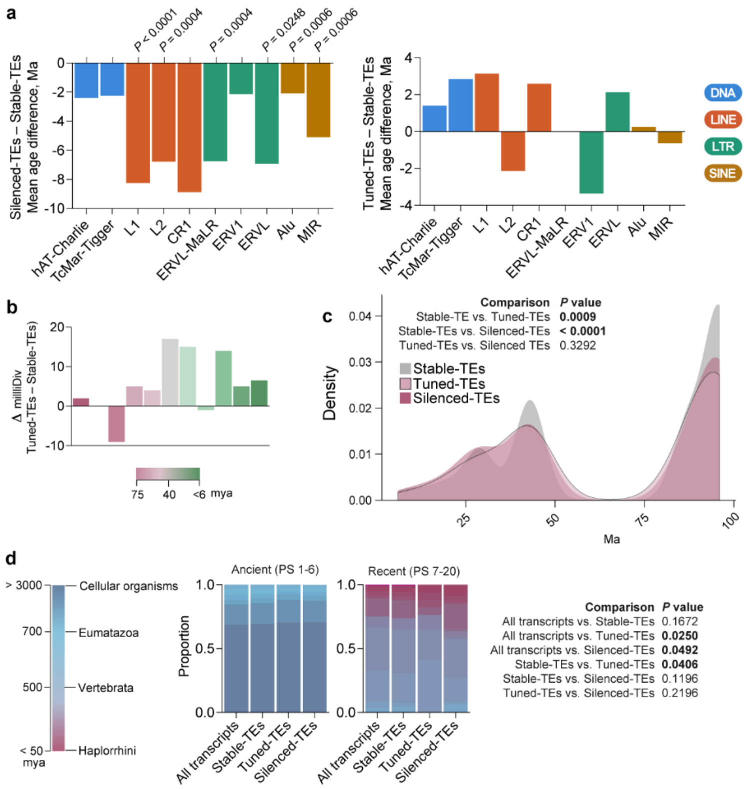


**Figure S4 | Younger and more mutated TE integrations are regulated by NMD and target genes of all ages. a**, Mean age difference in million years (Ma) between TE families in Stable-TEs and Silenced-TEs (left), and between Stable-TEs and Tuned-TEs (right). *P* values were calculated with a two-sided Wilcoxon rank-sum (Mann-Whitney U) test with BH correction. **b**, The difference in milliDiv values between Tuned-TEs and Stable-TEs for the most diverged TE milliDiv per transcript (in case of overlapping multiple TEs). *P* values were calculated within each age bin using a two-sided Wilcoxon rank-sum (Mann-Whitney U) test. **c**, Density distribution of the age of the oldest TE-flanking splice site in TE transcripts by the most distant species in which the splice site was conserved, excluding splice sites where this was human-specific. The *P* values were calculated by a two-sided Wilcoxon rank-sum (Mann-Whitney U) test, and included human-specific splice sites. **d**, Phylomapr age estimate of genes expressed by HeLa.control (All transcripts), or in the three TE groups (Stable-TEs, Tuned-TEs, Silenced-TEs). *P* values were calculated with Wilcoxon rank-sum (Mann-Whitney U) tests with BH correction.

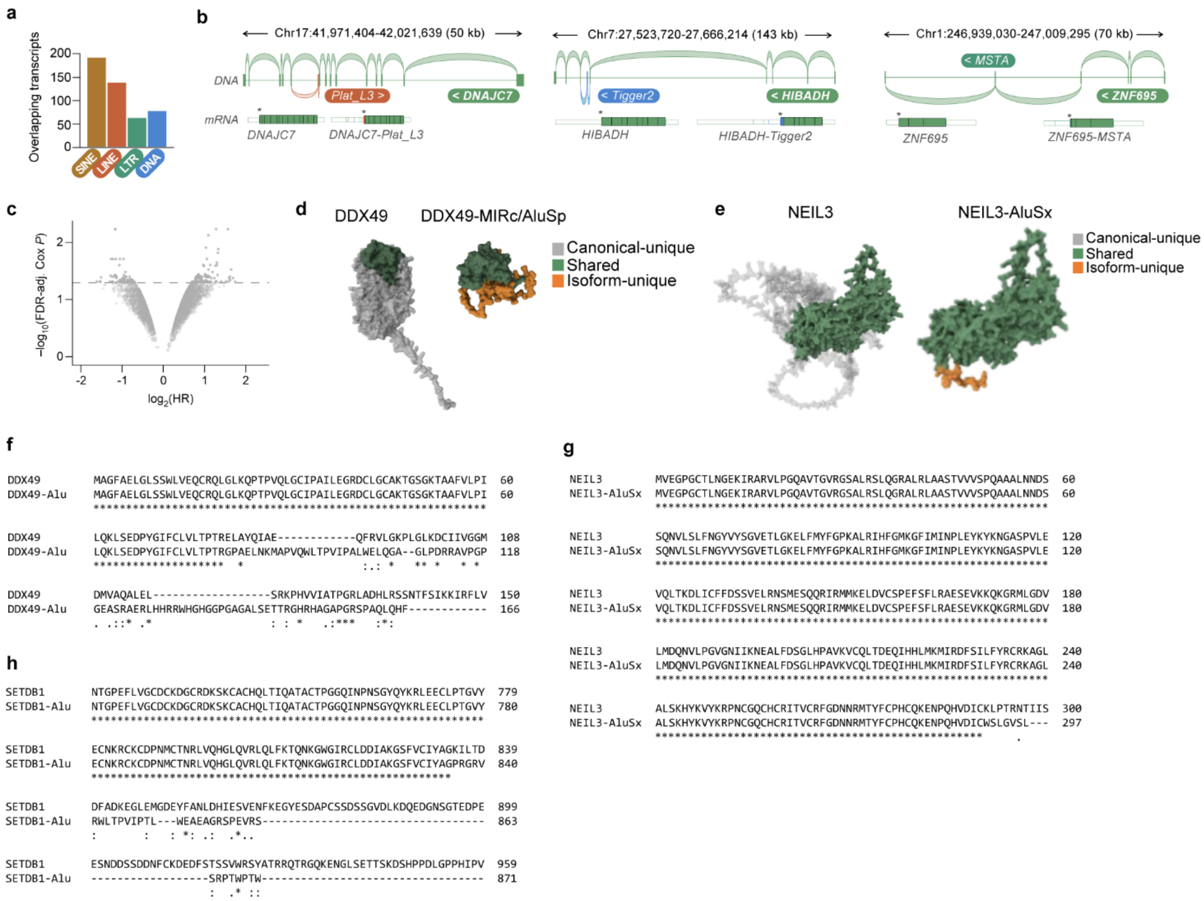


**Figure S5 | NMD regulates transcripts from all TE classes, involving a variety of types of splicing events and protein consequences. a**, FDI transcript overlaps by TE class. Some transcripts overlapped multiple TE families and so were counted multiple times. **b**, A Plat_L3 (*LINE*)-exonizing isoform of DnaJ Heat Shock Protein Family (Hsp40) Member C7 (*DNAJC7*) and a Tigger2 (*DNA*)-exonizing isoform of 3-Hydroxyisobutyrate Dehydrogenase (*HIBADH*), both targeted by NMD due to PTC introduction by the TE exon causing a frameshift. An MSTA (*LTR*)-exonizing isoform of Zinc Finger Protein 695 (*ZNF695*), where a mid-exon splice from the canonical terminal exon to a downstream *LTR* element was associated with a splice to another downstream exon. In this case, the MSTA element provided the SA, stop codon and SD sequences, truncating the protein (due to loss of most of the canonical terminal exon), as well as introducing a downstream EJC after splicing to the terminal exon, causing NMD targeting. **c**, Per locus NMD ratio survival association analysis in LUSC patients using an optimal NMD-ratio cutoff approach. Dashed line represents FDR-adjusted *P* value calculated by Cox (Wald) test. **d-e**, AlphaFold predictions of the protein structures of DDX49 and DDX49-MIRc/AluSp (d), and NEIL3 and NEIL3-AluSx (e), with coloring to indicate which approximate regions are shared or unique between isoforms. **f-h**, Amino acid sequence alignments of the entire proteins for DDX49 and DDX49-Alu (f), the entire proteins for NEIL3 and NEIL3-AluSx (g), and the truncated protein of SETDB1-Alu compared to SETDB1 (h). * indicates a match, : indicates strong conservation (different residues but similar properties), and . indicates weak conservation (slightly more different properties but still similar).

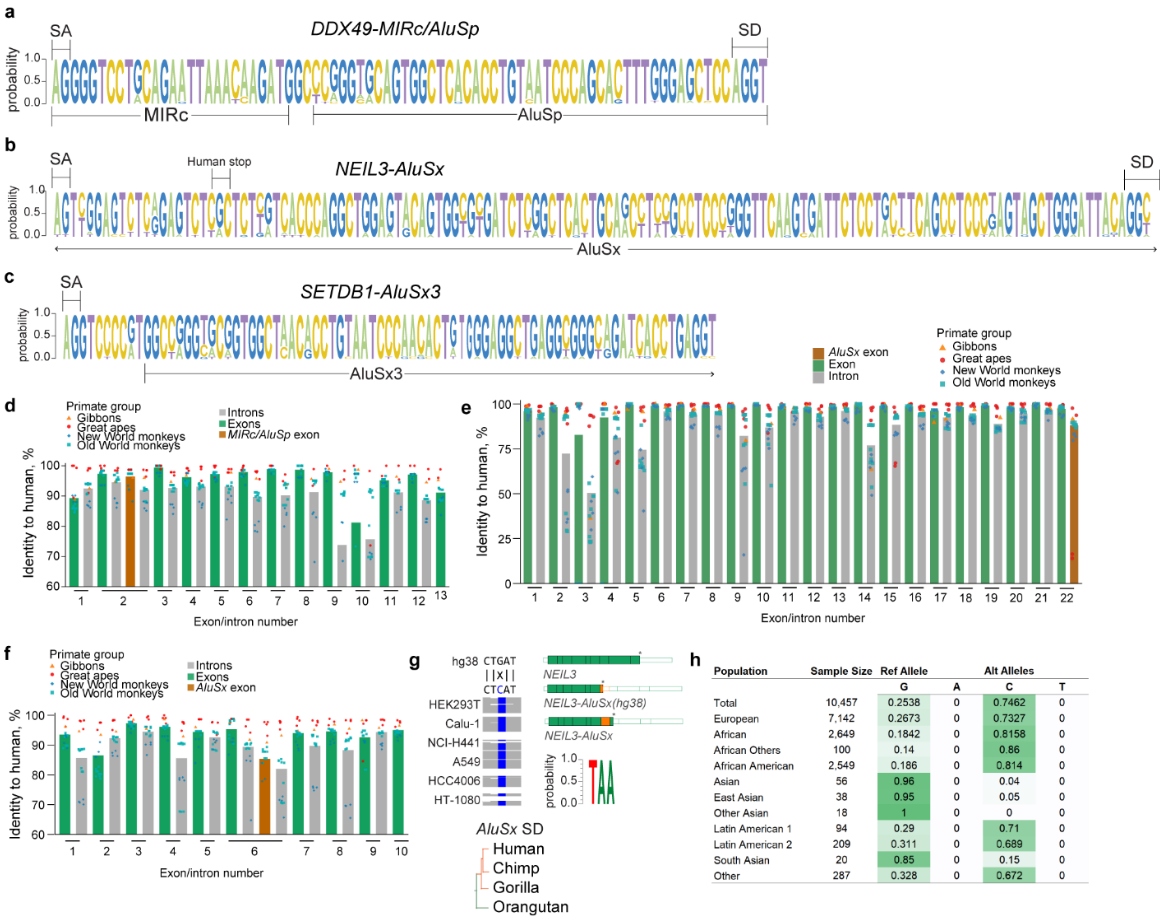


| Population | Sample Size | Ref Allele G | Alt Alleles A | C | T |
|---|---|---|---|---|---|
| Total | 10,457 | 0.2538 | 0 | 0.7462 | 0 |
| European | 7,142 | 0.2673 | 0 | 0.7327 | 0 |
| African | 2,649 | 0.1842 | 0 | 0.8158 | 0 |
| African Others | 100 | 0.14 | 0 | 0.86 | 0 |
| African American | 2,549 | 0.186 | 0 | 0.814 | 0 |
| Asian | 56 | 0.96 | 0 | 0.04 | 0 |
| East Asian | 38 | 0.95 | 0 | 0.05 | 0 |
| Other Asian | 18 | 1 | 0 | 0 | 0 |
| Latin American 1 | 94 | 0.29 | 0 | 0.71 | 0 |
| Latin American 2 | 209 | 0.311 | 0 | 0.689 | 0 |
| South Asian | 20 | 0.85 | 0 | 0.15 | 0 |
| Other | 287 | 0.328 | 0 | 0.672 | 0 |

**Figure S6 | The three *Alu*-overlapping NMD transcripts represent different levels of evolutionary conservation. a-c**, Logo plots for the splice acceptor (SA), splice donor (SD), and TE-overlapping regions for the novel exons present in *DDX49-MIRc/AluSp* (a), *NEIL3-AluSx* (b), and *SETDB1-AluSx3* (c) based on sequence conservation in 19 primate genomes. **d-f**, Mean percent identity of *DDX49* (d) *SETDB1* (e), and *NEIL3* (f) exons and introns across 19 primates, generated by a per-base gap-aware calculation. Symbols represent individual species genomes and are colored by primate group. Columns are colored by exons or introns, with the *Alu*-overlapping exon colored in brown. **g**, Schematic of the C-to-G mutation present in hg38 that was not sequenced in the tested cell lines, along with a schematic of the hg38 vs cell line-version of the *NEIL3-AluSx* transcript, a logo plot of the stop codon conservation across 19 primate genomes, and the point in evolutionary time when the *AluSx* SD site emerged (orange). **h**, Frequencies of the hg38 reference and alternative alleles in populations of different genetic ancestry for the rs6823747 SNP in *NEIL3-AluSx*. Data were obtained from the National Center for Biotechnology Information (NCBI) SNP database (https://www.ncbi.nlm.nih.gov/snp).

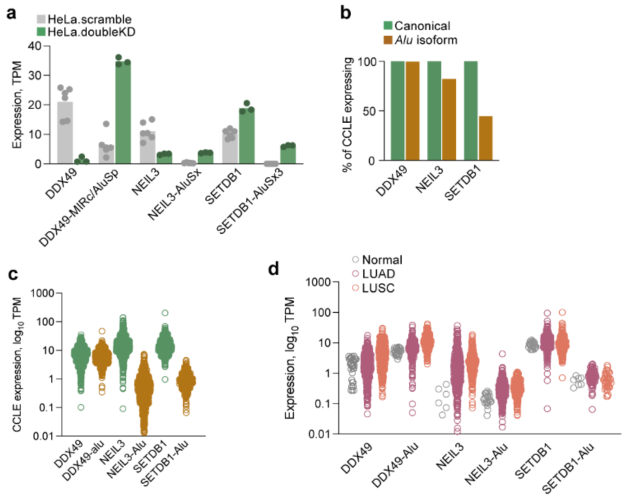


**Figure S7 | NMD-regulated *Alu*-overlapping isoforms are induced by NMD-blockade, and expressed across the CCLE and in Normal lung, and LUAD, LUSC cohorts. a**, Expression of canonical and *Alu*-overlapping isoforms of *DDX49*, *NEIL3*, and *SETDB1* in HeLa cell KD dataset[28,29] by aligning to NET assembly. **b-c**, Proportion of CCLE that expresses the same transcripts as in (a) after aligning to NET assembly (cutoff 0.1 TPM) (b), and transcript expression level (c). Symbols represent individual cell lines. **d**, Expression level of the same transcripts as in (a) in Normal lung, LUAD, and LUSC cohorts after realignment to NET assembly.

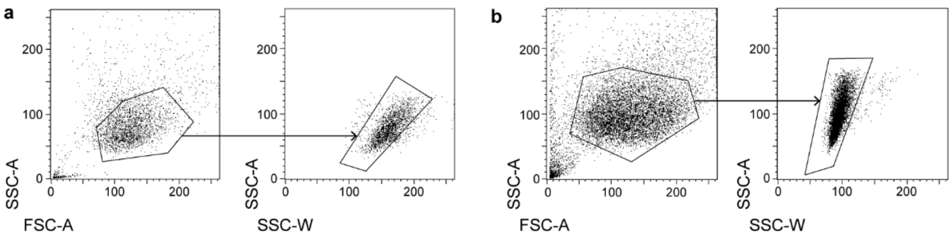


**Figure S8 | Flow cytometry gating strategies. a-b**, Representative flow cytometry plots showing gating strategies for experiments shown in Fig. 6a (a) and Fig. 6c (b).
